# Beyond the tripartite genome: a widespread, mobile and gene-rich accessory plasmidome expands the genomic landscape of the model symbiont *Sinorhizobium meliloti*

**DOI:** 10.64898/2026.11.19.767906

**Authors:** Juan José Picón-Cossio, Samantha G. Crossman, Andrew J. Price, Sean McGrath, Tania Gupta, Barney Geddes, Maria Alejandra Gil-Polo, Kayla Clouse, Sohini Guha, Liana Burghardt, Siva Sankari

## Abstract

*Sinorhizobium meliloti* is one of the best-studied nitrogen-fixing symbionts, yet research has focused almost entirely on its chromosome and two megaplasmids, pSymA and pSymB. Many strains carry additional accessory plasmids, but these large, pSymA-like elements are rarely recovered from short-read assemblies, leaving a hidden part of the genome unexplored. By sequencing 346 strains with long reads, we recovered 386 accessory plasmids from 76% of strains, with 318 absent from public databases, showing that accessory plasmids are the rule and not the exception. These plasmids were large (34–749 kb), including the largest accessory plasmid reported in S. meliloti, and most were mobilizable or conjugative (relaxases in 85% and mating-pair formation systems in 75% of plasmid clusters), equipping them to move between strains. RepA typing defined 13 replicon groups unique to *Sinorhizobium*, which could serve as a framework for tracking these plasmids in natural populations. We also found evidence of plasmid expansion by fusion, and one ∼315 kb plasmid was a cointegrate of two smaller plasmids from our collection. About two-thirds of plasmid gene families were absent from the main replicons, including defense systems, the anti-restriction protein ArdC, and biosynthetic gene clusters, 53% of which had never been observed before. Overturning the view that S. meliloti lacks restriction–modification systems, we found them in 153 genomes, including on 32 accessory plasmids. Most inferred gene transfers stayed within the family *Rhizobiaceae*. Accessory plasmids thus form a large, mobile and largely unexplored gene pool that may shape how this model symbiont competes and adapts.

## Introduction

*Sinorhizobium meliloti* is an economically important, experimentally tractable, gram-negative alphaproteobacterium from the family *Rhizobiaceae*. It exists as a free-living bacterium in a variety of soil environments and as an endosymbiont of leguminous plants in the genera *Medicago, Melilotus*, and *Trigonella* (1), where it fixes atmospheric nitrogen into plant-available forms. *S. meliloti* has long served as a model for bacterial endosymbiosis and biological nitrogen fixation processes. In particular, strains Rm1021 and Rm2011 have been used for laboratory studies for decades, and a multitude of genetic engineering techniques have been developed to study them. However, both strains harbor only the chromosome and two megaplasmids, leaving the accessory plasmids found in many other S. meliloti strains largely unexplored (2–4).

*Sinorhizobium meliloti* harbor a multipartite genome with a 3.65 Mb chromosome and two megaplasmids: a 1.35 Mb megaplasmid, pSymA, and a 1.68 Mb megaplasmid, pSymB (2, 5). pSymA primarily supports symbiosis, carrying genes involved in nodulation (*nod*) and nitrogen fixation (*nif, fix*) (6). In contrast, pSymB contributes to symbiosis through functions like exopolysaccharide biosynthesis, but is primarily associated with free-living fitness, encoding solute transport systems, transcriptional regulators, cell protection, and catabolic functions (7–9).

Because these three replicons are present in every strain and together encode the core of the symbiotic lifestyle, decades of research have centered on them, mostly in reference strains and a few wild isolates. Yet many *S. meliloti* strains carry additional plasmids. Unlike the stably maintained megaplas-mids, these vary from strain to strain and are termed cryptic or accessory plasmids—”cryptic” reflecting their elusive nature and our limited understanding of their functions (10– 16). However, a few studies suggest they carry genes that profoundly affect the fitness of the host, both in positive and negative ways (17, 18).

Plasmids in soil bacteria are associated with a variety of traits important for agriculture, medical, and biotechnological applications. They encode novel genes responsible for antimicrobial resistance (AMRs), Biosynthetic Gene Clusters (BGCs), Antimicrobial Peptides (AMPs), and defense systems (19). However, despite their functional relevance, plasmids from soil bacteria, especially rhizobia, are poorly characterized. This is partly due to their large size, often exceeding 100kb (19), and mosaic gene content, which together hinder their recovery from short-read assemblies (20, 21). In *S. meliloti*, accessory plasmids face the same limitations, which are further aggravated by their shared similarity with megaplasmids, especially pSymA (22).

The dynamics and diverse gene content of accessory plas-mids make them a rich reservoir for discovering new compounds with biotechnological relevance. There is also a growing clinical interest, as soil plasmids have gained attention for their potential to spread antibiotic resistance genes among clinically relevant bacterial species (19, 23, 24). Thus, comprehensive characterization and tracking of the species’ plasmids, also known as the plasmidome, is essential. Few studies have detailed the plasmidome of other rhizobial species such as *Rhizobium leguminosarum*, a clover-associated species characterized by harboring multiple large plasmids. These studies have identified highly diverse, gene-rich accessory plasmids involved in horizontal gene transfer (HGT) events. Some of these plasmids have been linked to declining partner quality under nitrogen fertilization, showing that changes in plasmid gene content can drive rapid evolution of symbiotic traits (25, 26).

Despite decades of research on *S. meliloti* as a model organism, only a few studies have reported the sequences of accessory plasmids from different *S. meliloti* strains (27–30). To address this gap, we assembled 346 S. meliloti genomes collected from different sources across the United States, Spain, France and Corsica using long-read sequencing (Oxford Nanopore or PacBio HiFi), from which we recovered 386 accessory plasmids. Together with 37 reference plasmids from public databases, we functionally annotated 423 accessory plasmids. These data reveal that accessory plasmids are present in approximately 76% of *S. meliloti* strains, establishing them as a standard component of the species’ genome rather than a peripheral feature. This represents a significant expansion of genomic data and provides a valuable resource for further investigation of the role of these extrachromosomal elements.

## Materials and Methods

### Strain Collection

MAG strains (n=39) were obtained from a previous collection (31). The AVT strains (n = 117) were isolated from nodules of three alfalfa varieties grown within 5 m of each other in an alfalfa variety trial at the Russell E. Larson Agricultural Research Center, Rock Springs, PA, USA. Plants were inoculated once at planting in 2016, and nodules were collected for the strain collection in fall 2020 (32).

The TG strains (n=184) were collected from alfalfa fields or pastures from 22 sites in North Dakota and one site in South Dakota. These included M. sativa (18 sites), M. falcata (2 sites), and Melilotus officinalis (3 sites). The bleached nodules were rinsed three times with Tryptone Yeast Extract (TY) broth, then crushed with a sterile pestle and streaked onto TY agar supplemented with 5 mM CaCl_2_. Plates were incubated at 28 °C for 3 days, and putative single rhizobia colonies were streak-purified twice. The purified strains were stored at *™*80 °C in 20% glycerol (v/v).

### Genome Library Preparation and Sequencing

MAG strains were sequenced at Stowers Institute for Medical Research: multiplexed PCR-free libraries (LongPlex Multiplexing Kit, seqWell; SMRTbell Prep Kit 3.0, PacBio) were size-selected (*≥* 10 kb), pooled with non-size-selected library at a 5:1 molar ratio and sequenced on a PacBio Revio. AVT strains were sequenced on a PacBio Sequel IIe at the Penn State Huck Institutes of the Life Sciences Genomics Core Facility (32). TG strains were sequenced by Plasmidsaurus (Eugene, OR, USA) with amplification-free Oxford Nanopore (ONT) V14 chemistry on a PromethION with R10.4.1 flow cells. Details for MAG and TG strains are given in Supplementary Methods S1.1.

### Genome Assembly, Quality and Taxonomy

HiFi reads were filtered with fastp-long v0.4.1 (33). HiFi and Oxford Nanopore (ONT) reads were assembled with Flye v2.9.6 (34), and ONT assemblies were polished with Medaka v2.1.1. Assembly quality and completeness were assessed with QUAST v5.3.0 (35) and CheckM (36), and contigs were assigned to the chromosome, pSymA, pSymB or “Other” by alignment to a reference genome with minimap2 v2.28 (37); Supplementary Methods S1.2). Species assignment was confirmed by a Mash-distance phylogeny of 3,308 Rhizobiaceae RefSeq genomes built with Mashtree v1.4.6 (38). Average nucleotide identity (ANI) pairwise between strains and to *S. meliloti* Rm1021 (GCF_000006965.1) were computed with fastANI v1.34 (39) using default parameters.

### Accessory Plasmid Identification and Typing

Contigs shorter than 1 Mb were pooled with *S. meliloti*-related plas-mids from PLSDB 2024v2 (40) and clustered by sequence containment with MobMess (41), which classifies clusters as backbone, compound, maximal or fragment. Fragment clusters were discarded, and only plasmid-s ized clusters were retained (Supplementary Methods S1.3). MobMess clusters (*≥* 90% coverage and identity) were used as plasmid units in all downstream analyses, since they dereplicate the dataset without over-merging genuinely distinct plasmids. Retained sequences were classified as accessory plasmids if Bakta v1.12.0 (42) annotated a replication or partition protein or geNomad v1.12 (43) detected a relaxase, and low-depth sequences were curated manually (Supplementary Methods S1.3). The final set comprised 423 accessory plasmids: 386 from the strains sequenced here and 37 reference plasmids from PLSDB.

Plasmid 3-mer composition was compared using Bray–Curtis distances and UMAP (44), and ANI and alignment fraction (AF) were computed with skani v0.3.2 (45). Relaxase (MOB) and mating-pair formation (MPF) types and nearest Mash neighbors were assigned with MOB-typer v3.1.9 (46). Replicon groups were defined by clustering one representative RepA per plasmid, taken preferably from a complete repABC operon, with the R. leguminosarum Rh group references of Cavassim *et al*. (25) and Vereau Gorbitz et al. (26) using MMseqs2 v18 (Supplementary Methods S1.3).

Plasmid copy number was estimated as the median read depth of each plasmid divided by that of the chromosome in the same assembly, after mapping reads with minimap2 v2.28 and computing per-base depth with SAMtools v1.11 (47); pSymA and pSymB served as single-copy controls (Supplementary Methods S1.3). GC content and length of each main replicon were compared with those of accessory plasmids using two-sided Mann–Whitney U tests.

The presence of accessory plasmids was verified in 21 strains by a modified Eckhardt gel electrophoresis protocol (see (48)).

### Functional Annotation

Plasmids were annotated with Bakta v1.12.0 (42). Toxin–antitoxin (TA) genes were identified with TASmania HMMs (49), biosynthetic gene clusters (BGCs) with antiSMASH v8.0 (50), defense and anti-defense systems with DefenseFinder v2.0.1 (51), and CAZymes with run_dbcan (52), keeping CAZymes supported by at least two dbCAN tools; signal peptides were predicted with SignalP 6.0 (53). Following Pulliam et al. (54), BGCs were grouped into gene cluster families (GCFs) with BiG-SCAPE v2.0.3 (55) and BiG-SLiCE v2.0 (56, 57). A BGC was considered known if its BiG-SCAPE GCF (cutoff 0.3) contained a MIBiG 4.0 reference, reported if it lay within a BiG-SLiCE distance of 0.65 of an antiSMASH-DB v4 GCF (i.e., previously observed in public genomes), and novel otherwise (Supplementary Methods S1.4).

### Gene Content Analysis

A pangenome of all replicons was built with Panaroo v1.6.0 (58), and its gene families were used to quantify gene sharing between accessory plasmids and the main replicons. COG categories were assigned with eggNOG-mapper v2.1.13 (59), and COG enrichment per plasmid cluster was tested with Fisher’s exact test and Benjamini–Hochberg correction (Supplementary Methods S1.5).

### Horizontal Gene Transfer Events

Putative horizontal gene transfer (HGT) was inferred from the weighted gene repertoire relatedness (wGRR; (60)) between the 423 accessory plasmids and PLSDB plasmids *≤* 500 kb that aligned to at least one of them. Plasmid pairs with 0.001 *<* wGRR *<* 0.1 sharing at least one bidirectional best hit with *>* 80% identitywere considered putative HGT connections (Supplementary Methods S1.6).

## Results

### Genome Assembly Quality and Strain Taxonomic Identity

A total of 346 *Sinorhizobium meliloti* genomes were assembled from three long-read datasets: in-house PacBio HiFi (MAG), PacBio HiFi from Penn State (AVT), and Oxford Nanopore from Dakota State (TG). All 346 genome assemblies were of high quality, with completeness ranging from 98.49% - 99.59% (median 99.56%; **Fig. 1A**). Assemblies were highly contiguous, with a median of 5 contigs per genome (range 2–14; n = 341): 4 for the MAG, 6 for the AVT, and 5 for the RG.

**Fig. 1.**
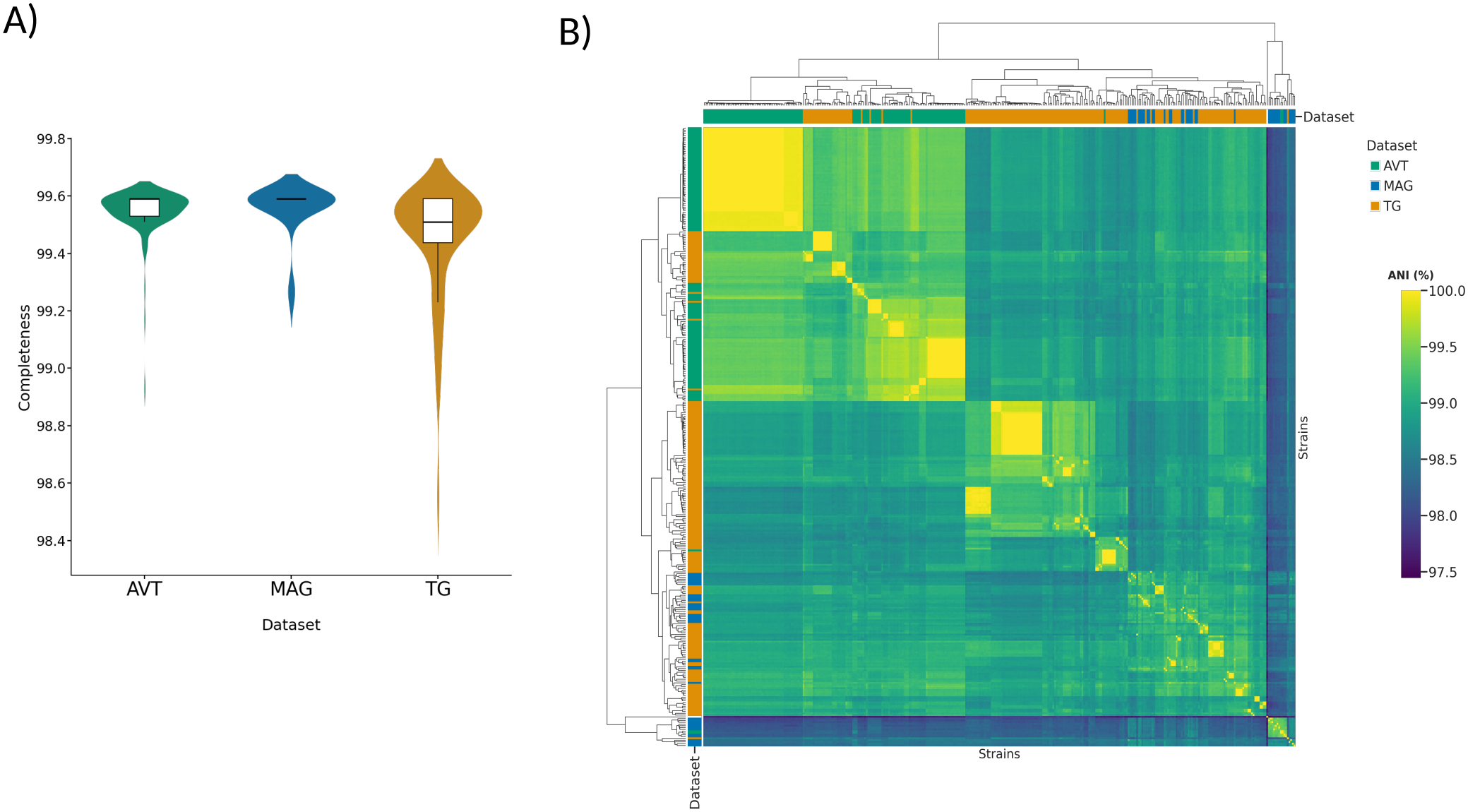
**A)** Percentage completeness according to CheckM. **B)** strains all vs all ANI comparison

MAG genomes contained the least number of contigs (median 4, range 3–6), followed by TG (median 5, range 2–14) and AVT genomes (median 6, range 3–11). These differences were mostly attributable to data coverage (median 45x; **Fig. S1A**). Each contig was classified as Chromosome, pSymA, pSymB or ‘Other’, the last containing accessory plasmids, fragments or artifacts. Fragmentation occurred mainly in the megaplasmids and accessory plasmids. The median GC content of ‘Other’ contigs was lower than that of pSymA, which in turn was lower than that of pSymB and the chromosome in all the datasets (**Fig. S1A)**. Most assemblies presented at least three replicons greater than 1 Mb, representing the chromosome, pSymA, and pSymB (**Fig. S1A)**.

All strains were placed within *Sinorhizobium*, and all genomes shared 97.8–99.97% average nucleotide identity (ANI) with the reference strain Rm1021 (median 99.05%), confirming their assignment to *S. meliloti*. Pairwise average nucleotide identity (ANI) exceeded 99%. The AVT dataset showed the highest in-group similarity, followed by TG, while MAG strains displayed the broadest spread (**Fig. 1B**). This pattern is consistent with sample origin: AVT strains were collected from a single agricultural forage field at a single timepoint, TG strains from a wild source, and MAG strains are a subset of the wild isolates collected in Spain, France, and Corsica by Riley et al. (31).

### Accessory Plasmid Identification and Characterization: A Containment-Based De-Replication Approach

Because sequence similarity with pSymA can leave accessory plasmids fragmented, we de-replicated contigs by sequence containment with MobMess, which separates complete plasmids from fragments (**Fig. S2A-C**). Excluding fragments, we identified 423 accessory plasmids in 148 clusters, 386 from our strains and 37 from PLSDB. Of the 386 plasmids recovered here, 318 (119 clusters) had no database counterpart. Very few clusters were shared between datasets (only 5 between TG and AVT), suggesting ecological or geographic specificity. TG produced the largest number of clusters (81 from 220 sequences); AVT, despite containing 133 sequences, yielded only 8 clusters, consistent with its high within-collection strain similarity; and MAG, the most diverse by ANI, produced 28 clusters from 33 sequences (**Fig. S2D**).

Differences in evolutionary dynamics among megaplasmids, such as those distinguishing pSymA from pSymB, can be reflected in features such as GC content and codon usage. We therefore first characterized contigs using Bray–Curtis distances of trinucleotide (3-mer) composition. This separated the chromosome, pSymA, pSymB and accessory plasmids (**Fig. 2A**), with the exception of two accessory plasmids that clustered with pSymA. Accessory plasmids therefore generally carry compositional signatures distinct from those of the three main replicons, in line with earlier dinucleotide-signature analyses in this species (61). Accessory plasmids had a median GC content of 58.97 (range 57–61%), lower than that of pSymA (median 60.17%), pSymB (62.39%) and the chromosome (62.67%), and sizes of 34–749 kb (**Fig. 2B and 2C**). To our knowledge, the 749 kb plasmid identified here is the largest accessory plasmid reported in this species. The smallest accessory plasmid was a single 7.5 kb element harboring ten hypothetical proteins, whose nearest Mash neighbor was pRm1132f (NC_004965), a 7.2 kb *S. meliloti* accessory plasmid first described by Barran *et al*. (62).

**Fig. 2.**
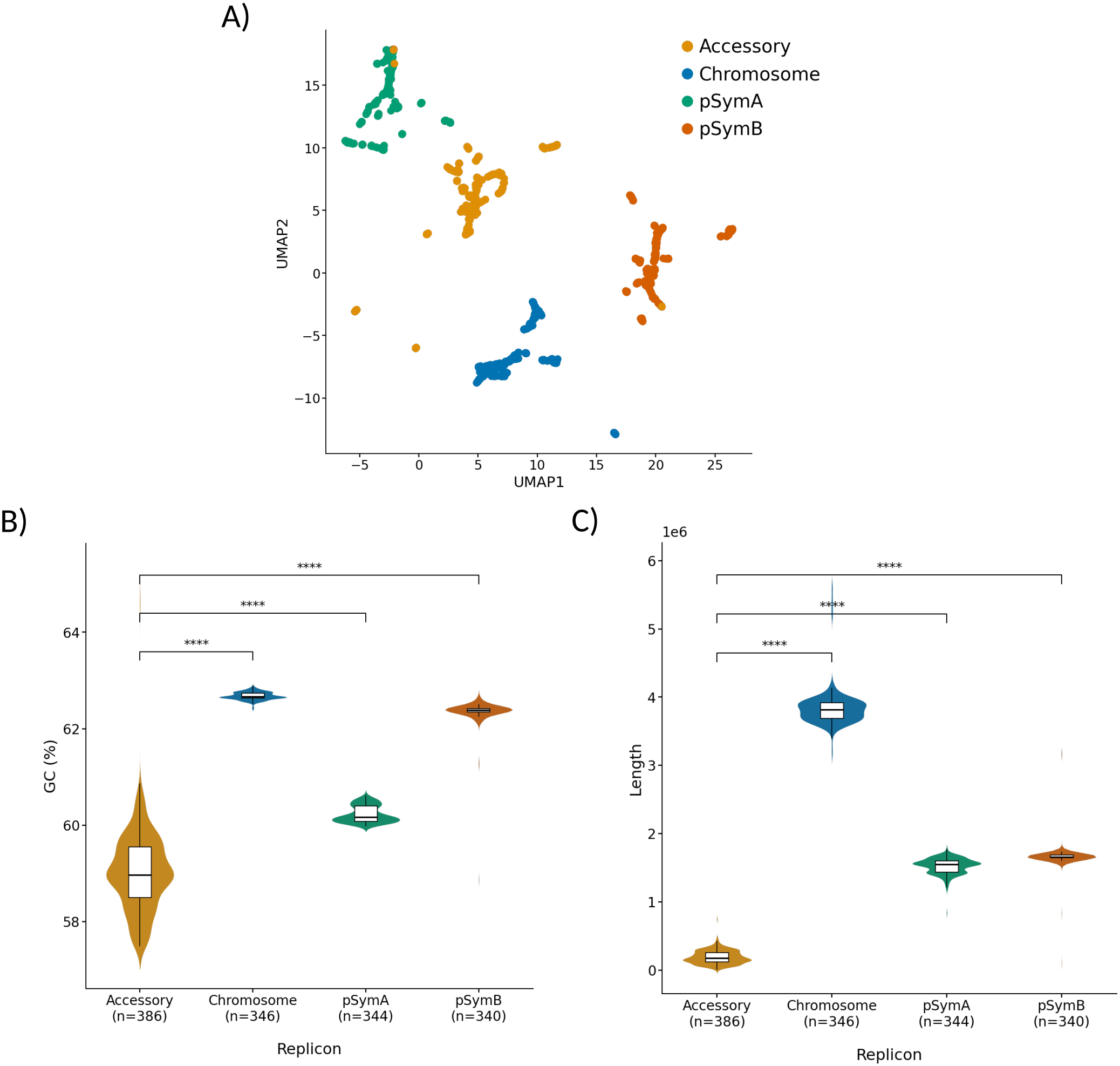
**A)** Trinucleotide (3-mer) distances projected on a UMAP. Each dot represent a sequence. Colored by replicon. **B)** GC content of Chromosome, pSymA, pSymB and accessory plasmids. **C)** Length of Chromosome, pSymA, pSymB and accessory plasmids Each main replicon was compared with accessory plasmids using a two-sided Mann–Whitney U test.

In total, 258 of the 346 strains harbored at least one accessory plasmid (**Fig. S2E**). Based on sequencing coverage relative to that of the chromosome, most plasmids were estimated to be present at approximately one copy or fewer per cell. Some plasmid clusters fell well below one, which may reflect plasmid loss during isolate culture, heterogeneous subpopulations within an isolate, or technical effects such as uneven coverage. On the other hand, some accessory plasmids were estimated to be present in *≥* 2 copies (**Fig. S2F and S2G**).

Accessory plasmids were confirmed in a subset of strains using a modified Eckhardt gel electrophoresis protocol (48). Representative gels are shown in **Fig. S2H**.

### Most Accessory Plasmids Have the Genetic Potential for Conjugation

Large plasmids (above ∼30–50 kb) tend to be mobilizable or conjugative, since conjugation becomes the most efficient route of horizontal transfer for plasmids of this size range (63). Moreover, conjugation has been extensively documented in rhizobia (16, 64). The *S. meliloti* plasmidome is dominated by large replicons, with 95% exceeding 70 kb. Consistent with this, 75% of clusters (313 sequences) carried a mating-pair formation (MPF) system (**Fig. 3A**), the type IV secretion machinery required for conjugative DNA transfer (65), indicating that conjugation is a major route of plasmid dissemination in this species as well (63, 66).

**Fig. 3.**
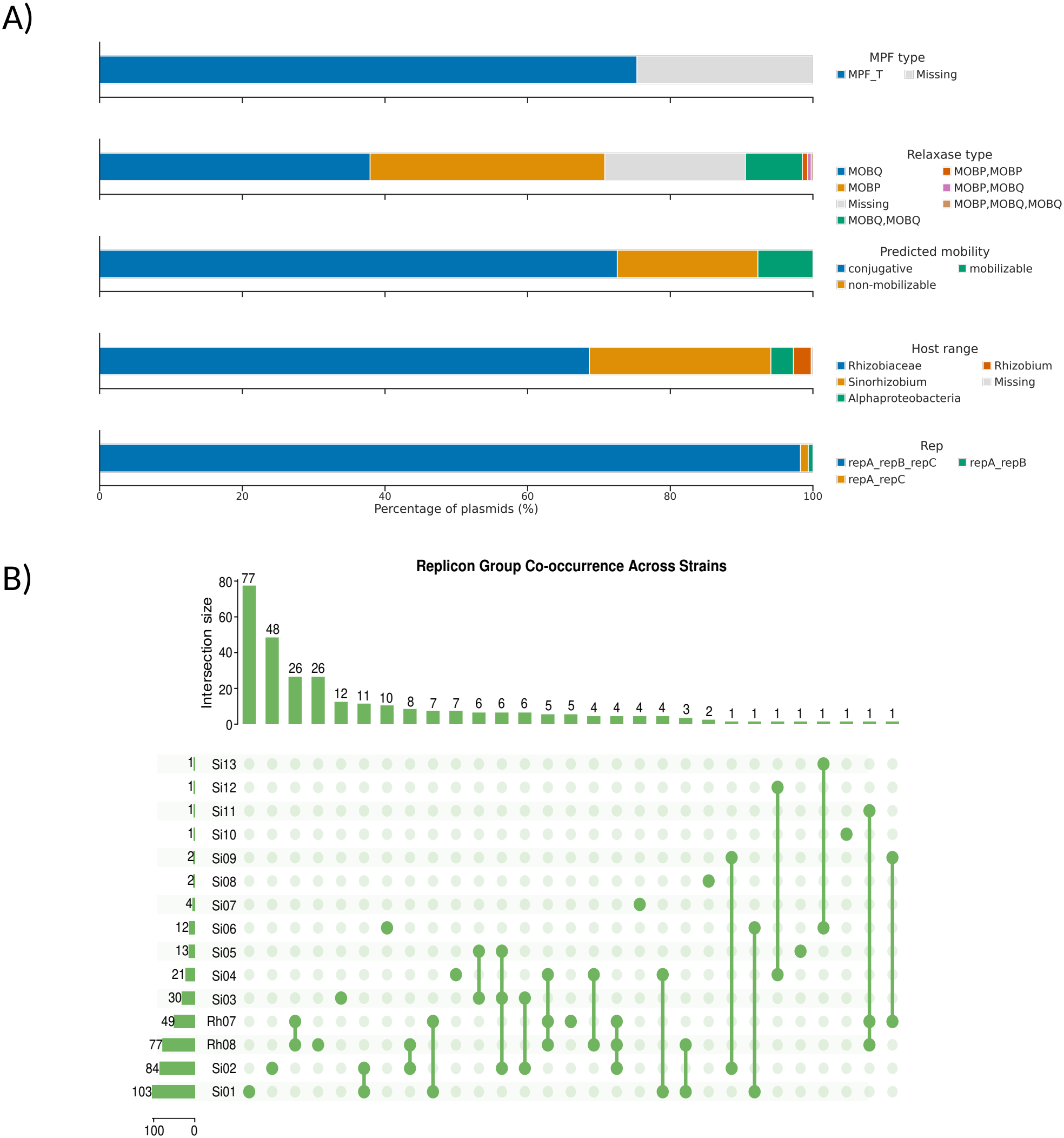
**A)** Percentage of Relaxases, MPF, and Replication protein annotations across accessory plasmids. Bars are colored by the type indicated in the right legend. **B)** Co-occurrence of replicon groups across strains. The top bar chart shows the number of strains. The left bar chart shows the number of plasmids. Connected dots in the matrix denote which replicon groups co-occur in each intersection

Relaxase, the protein that initiates and terminates plasmid mobilization (67), was present in 85% of accessory plas-mid clusters (338 accessory plasmids; **Fig. 3A**). MOBQ and MOBP were the dominant relaxase families, consistent with previous reports for Rhizobiaceae plasmids (68). The only exception is the 7.5 kb plasmid for which geNomad predicted a MOBV-type relaxase. Twelve clusters (39 sequences) carried multiple relaxases, in some cases from different families. For example, TG114 and TG34 carry a ∼420 kb circular plasmid with chromosome level coverage annotated with both MOBP and MOBQ relaxases. Plasmids carrying multiple relaxases may represent remnants of past cointegrates formed through recombination with other mobile genetic elements (MGEs).

Most accessory plasmids were predicted to be mobilizable or self-transmissible, whereas only 20% were classified as non-mobilizable, corresponding to 21 plasmid clusters. Regarding host range, most accessory plasmids were predicted to be restricted to the family level (*Rhizobiaceae*), with only a few showing a broader host range at the class level (**Fig. 3A**).

Most sequences carried replication proteins. Of the 423 accessory plasmids, 403 carried at least one repA gene and 394 carried a complete repABC operon (**Fig. 3A and S3A**). Twenty sequences lacked an explicit RepABC protein but carried other partition/replication annotations (ParB/RepB/Spo0J family protein, ParA), possibly indicating divergent replication modules. RepA was used to assess replicon group diversity, restricted to replicons carrying a single copy of the gene.

### Replicon Typing Defines Sinorhizobium-Specific Replicon Groups

Replicon-based classification informs plasmid incompatibility (69). In alphaproteobacteria this role is dominated by the repABC family, which is absent from the Enterobacteriaceae on which the canonical Inc scheme was built (70–72). We extracted and clustered repA from 401 accessory plasmids. Of these, 49 plasmids clustered with Rh07 and 77 with Rh08, suggesting possible ancestral plasmid exchange between *Rhizobium* and *Sinorhizobium*. The remaining 275 sequences formed 13 novel *Sinorhizobium*-specific clusters, designated Si01–Si13 in order of decreasing abundance (**Fig. 3B and S3B**), with 80% of plasmids concentrated between Rh07 and Si03. Most strains carried a single accessory plasmid, yet those with two or three carried plasmids from different replicon groups (**Fig. 3B and S2E**).

Pairwise ANI showed that plasmids were largely unique across most of their length: high-identity regions (*>* 80%) were concentrated at AF *≤* 40% (**Fig. S2D**), likely corresponding to shared backbones. Because replicon- and MOB-based schemes poorly capture such genome-wide similarity (41, 73), containment-based clusters provide a better unit for comparing plasmids, grouping near-identical plasmids without merging distinct ones that share only a backbone.

### Accessory Plasmids Harbor Type II and Type IV Toxin–Antitoxin Systems and Represent a Source of Novel Functions

Beyond replication, plasmids often encode maintenance systems that ensure their stable inheritance, including active partition and toxin–antitoxin (TA) systems. These are particularly important for low-copy-number plasmids, which cannot rely on random segregation to reach both daughter cells (74, 75).

TA genes were widespread on accessory plasmids. We annotated 2,071 TA genes (1,207 toxins and 864 antitoxins) on 413 of the 423 accessory plasmids or 141 of the 148 clusters. ParDE, the type IV AbiEii/AbiGii toxins, RHH, and type II Phd/YefM–Doc TA were the most frequent, present on 44–56% of plasmids (**Fig. 4A and 4B**). TA gene density decreases as plasmid size increases (**Fig. S4A**) consistent with previous reports and the proposal that, in larger replicons, TA systems are progressively displaced by other beneficial accessory genes (76, 77).

**Fig. 4.**
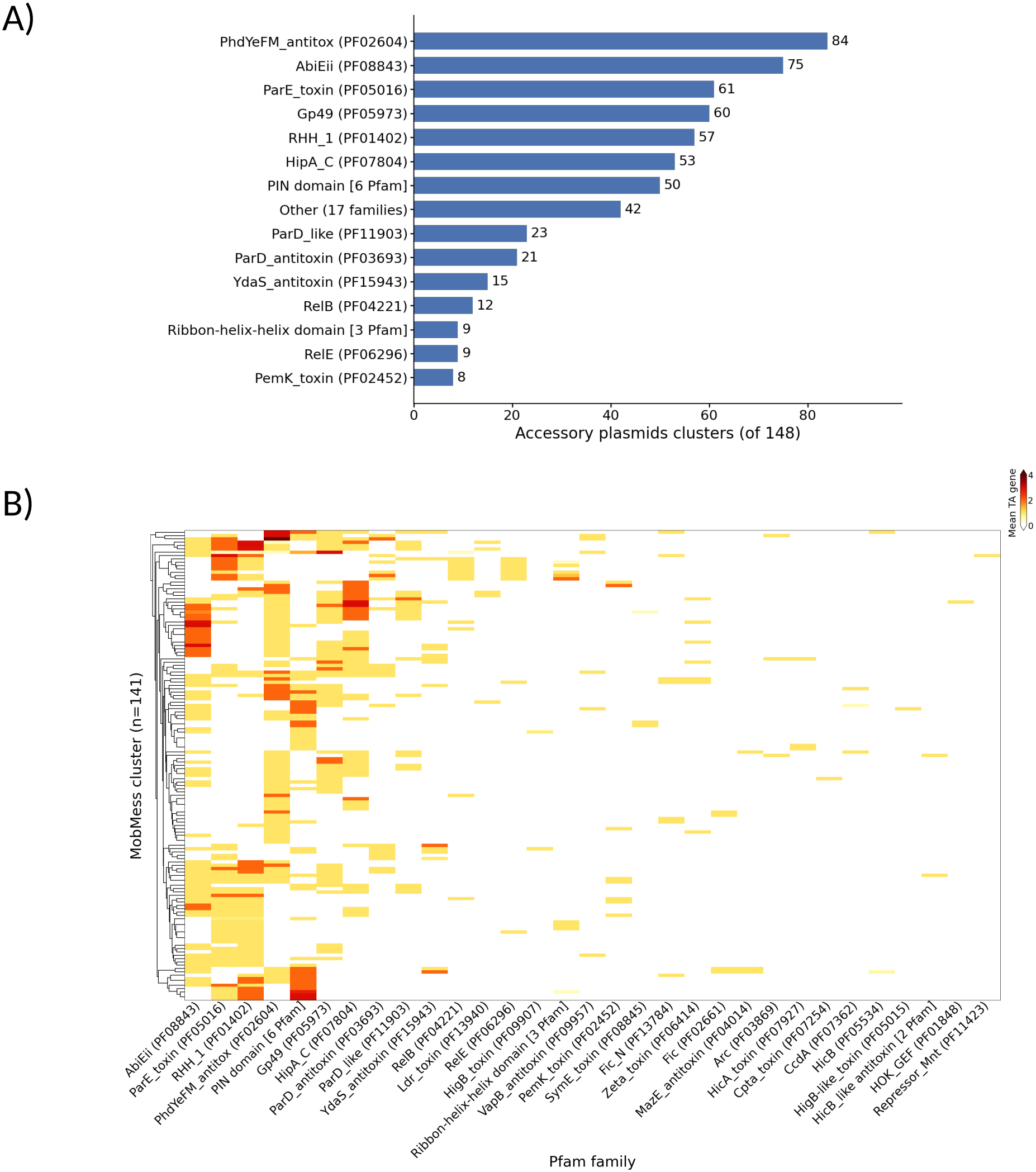
**A)** Percentage of accessory plasmids carrying at least one gene of each TA family. Families present on less than 5% of plasmids (MazEF/PemK, HicAB, zeta toxin, and CcdAB) are combined as “Other”. **B)** Mean number of TA genes for each family across accessory plasmid clusters. Rows are plasmids clusters and columns are Pfam families associated with TA HMMs.

Several predicted TA genes encode domains associated with ParB-like nucleases, PIN domains, transcriptional regulator families such as the CopG family, and unknown-function domains, making them candidates for novel TA-like activities. (**Fig. S4B**).

### Cointegration with MGEs Contributes to Large Accessory Plasmid Size

Cointegration between plasmids and other MGEs is an important driver of plasmid evolution. By merging two elements into a single replicon, cointegration diversifies gene content, circumvent incompatibility between co-resident plasmids, enables the transfer of non-conjugative elements across hosts, and can amplify antimicrobial resistance by assembling multidrug-resistant megaplasmids (78– 83). Some accessory plasmids of *S. meliloti* carried multiple copies of individual Rep proteins. **Fig. 5A** shows the number of annotations found for proteins related to the RepABC module as well as annotations related to replication or partition.

**Fig. 5.**
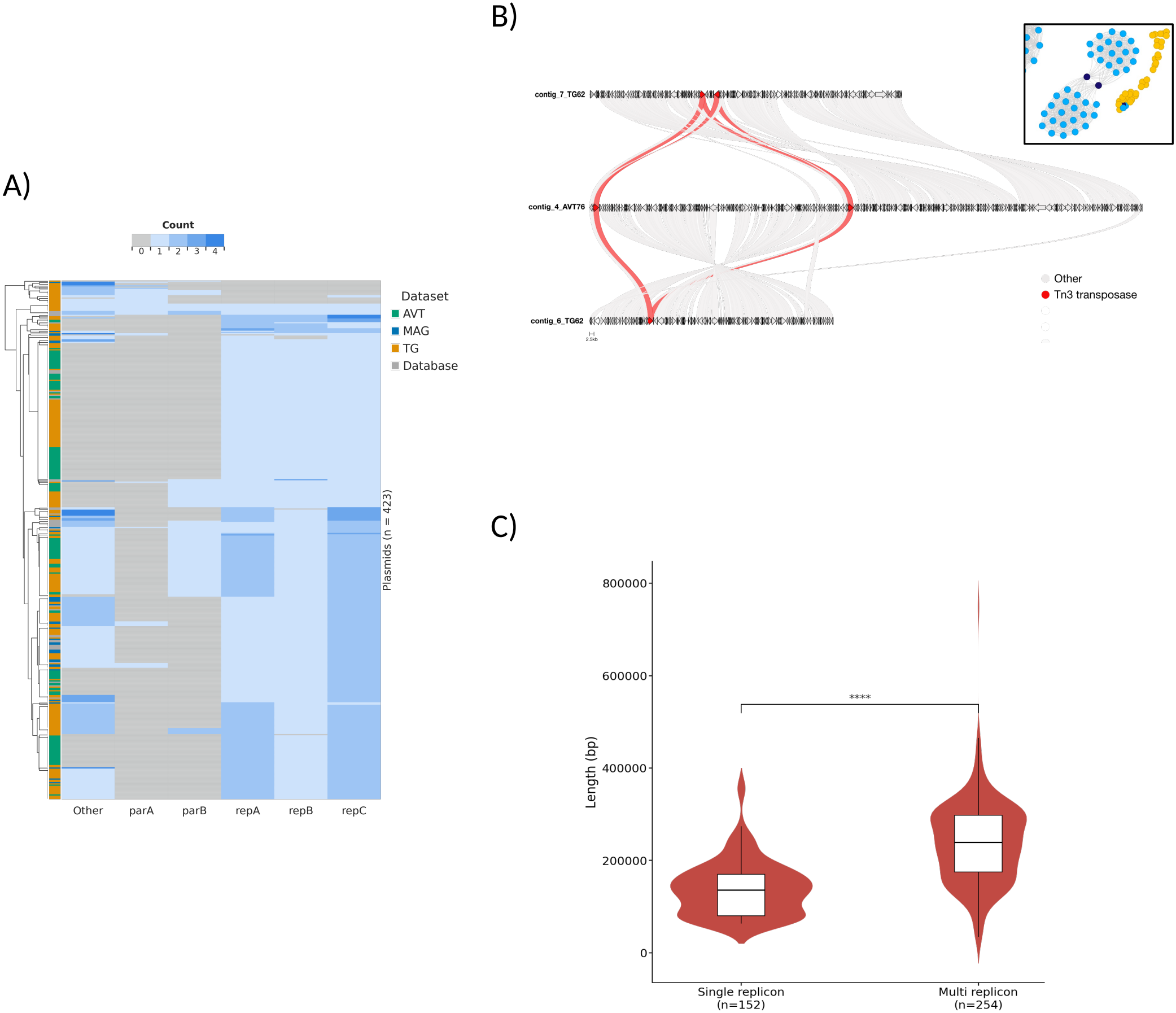
**A)** Number of replication proteins per plasmid. Rows are colored by dataset. Category Other means any other annotation related to a replication/partitioning protein. **B)** Protein alignment between a representative plasmid cointegrate (middle) and its parental plasmids (top and bottom). Red links highlight alignments between Tn3 transposases. Inset: the cointegrate visualized within the containment network; cointegrate nodes are shown in deep indigo and parental plasmid nodes in sky blue. **C)** Plasmid length by replicon multiplicity. A multi-replicon group is defined as sequences with at least one rep gene from the RepABC duplicated

Compound clusters in MobMess represent a way to investigate extra DNA sequences gained/lost by plasmids. Among our accessory plasmids, we found 19 compound clusters (37 plasmids) which, compared with their parent backbones, had acquired 77 regions with a median of 34 kb, most of them carrying transposases (67). These regions carried putatively adaptive cargo genes, including sulfur-acquisition cassettes (CysND, sulfotransferase, sulfatase, TauABCD), genes for phosphonate and polyamine metabolism, ectoine synthesis, aromatic-compound degradation, ACC deaminase and nonribosomal peptide synthetases (NRPS). They also included replication genes (23 regions), MOB/MPF modules (15), and ICE- or IME-like elements (6), while 33 carried none of these markers (**Fig. S5A and Supplementary data**). Replicons in *S. meliloti* carrying a second replication module have been reported (74) and may have formed by cointegration (84).

Interestingly, in one case we found entire repABC operons were present in two copies. Together with the size distribution skewed toward larger replicons, this led us to ask whether fusion contributes to the large size of these replicons. Cluster 381 (AVT76 and AVT109, ∼315 kb) is a cointegrate case for which both parents, PS43 (cluster 380; ∼158 kb) and PS41 (cluster 364; ∼146 kb), were sampled. Each parent covered about half of the compound replicon, and the two were joined by Tn3-family transposases (**Fig. 5B**), a transposase family previously observed during cointegrate formation (85). Although this was the only cointegrate for which both parents were sampled, it is unlikely to be the only one. In 7 clusters (9 plasmids), mostly built on the PS10 or PS33 backbones, acquired regions carried a replication gene with no matching second parent in the dataset. These are likely cointegrates whose second plasmid was not sampled (**Fig. S5A**).

Because detection by containment requires that the cointegrate and both of its parents persist in the dataset, it recovers only relatively recent events and cannot estimate how often fusion occurs. Nonetheless, larger plasmids tended to carry more copies of RepABC-associated proteins (**Fig. 5C and S5B**), consistent with fusion contributing to plasmid size, as has been reported in *Staphylococcus aureus* (81).

### Accessory Plasmids Carry Anti-Restriction Systems Distinct from pSymA and Restriction-Modification Systems

Restriction–modification (R-M) systems are the most widely distributed and studied bacterial defense mechanisms developed against invading DNA such as bacterio-phages, found in over 90% of bacterial genomes (86).

R-M systems are typically chromosomal, although they have also been found on mobile genetic elements such as plasmids (86, 87). *Sinorhizobium meliloti* has long been considered to lack R-M systems. The only one reported is a type I system in strain Rm1021, which carries just the three main replicons. It is flanked by insertion sequences, suggesting that it was acquired horizontally. (88–90). While annotating toxin–antitoxin systems, however, we noticed that many accessory plasmids encoded ArdC, an anti-restriction protein proposed to protect plasmids from host R-M systems during conjugation (91–93). We therefore asked whether *S. meliloti* acquires defense systems, including R-M, through accessory plasmids.

On the main replicons, defense systems were concentrated on the chromosome and pSymA, and pSymB carried them in only a few strains. The most common systems there were SoFIC (340), SanaTA (166), and Gabija (134) (**Fig. 6A**). On accessory plasmids, the most common were DS-20 (74), Gabija (62), and Lamassu-Fam (61) (**Fig. 6B**).

**Fig. 6.**
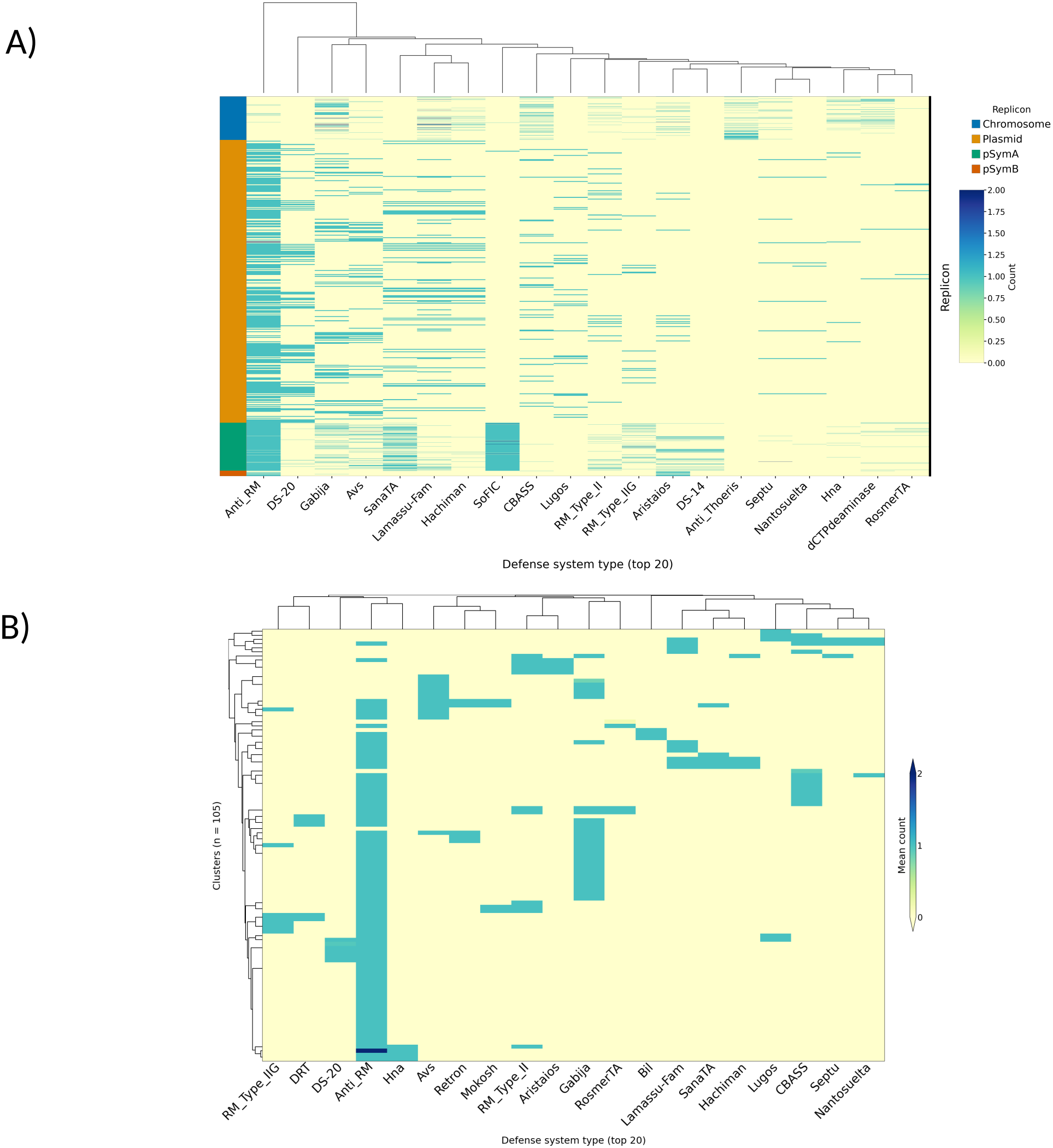
**A)** Heatmap showing the number of defense system types for chromosome, megaplasmids, and accessory plasmids. The top 20 defense system types are plotted. Rows represent a replicon sequence indicated by the row color. Columns represent defense system types. **B)** Heatmap showing the mean number of defense system types per plasmid cluster. The top 20 defense system types are plotted. Rows represent plasmid clusters, and columns represent defense system types

Contrary to the view that *S. meliloti* lacks R-M, 153 genomes carried at least one R-M system, most often type II (107 genomes) and less often type I (46). Accessory plasmids also contributed R-M systems, with type II systems in 18 clusters (32 plasmids). Two anti-restriction proteins were found between pSymA and accessory plasmids; ArdC, which protects plasmids against type II R-M during conjugation (91, 92), was annotated in 58% of accessory plasmid clusters (86 clusters; 232 plasmids). In contrast, ∼97% of pSymA replicons carried ArdB, which protects against type I R-M (94). The partitioning of ArdC and ArdB between accessory plas-mids and pSymA suggests that the two replicon classes have adapted to different R-M barriers during transfer.

### Accessory Plasmids Differ in Gene Composition, and They Share Genes Mostly with pSymA

Gene redundancy has been linked to functional innovation and environmental adaptation across bacteria (95–97), and it has been reported to be significantly high in *S. meliloti* (8). We inspected it using the gene families established with Panaroo.

About 34% of accessory-plasmid gene families were shared with at least one main replicon. Normalized by the number of gene families on each replicon, accessory plasmids shared the most with pSymA (29.6% of families), followed by the chromosome (12.1%) and pSymB (11.5%), suggesting that pSymA is the main partner of accessory plasmids in gene exchange (**Fig 7A)**, consistent with previous reports (22).

**Fig. 7.**
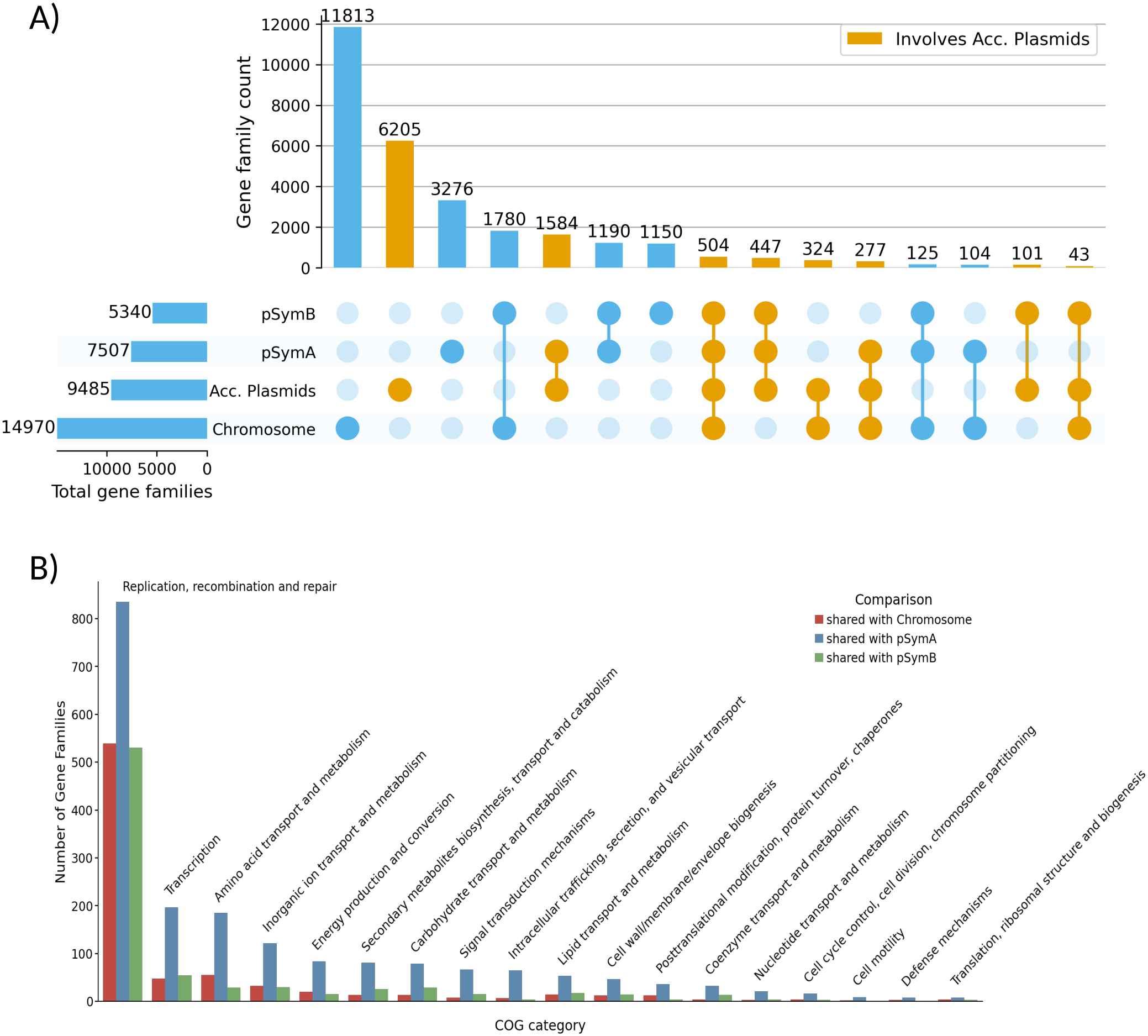
**A)** Number of genes shared between the chromosome, pSymA, pSymB, and accessory plasmids. Colored in orange when the genes are shared with accessory plasmids. **B)** Frequency of COG categories shared between accessory plasmids and the main replicons. Bars are colored by the comparison. The count value is the number of gene families per COG category in each comparison.

Families shared with the chromosome were mostly recombination-related, such as transposases (98), with a few in “Transcription” and “Amino acid transport and metabolism.” “Replication, recombination and repair” was also the most frequent category among families shared with pSymA and pSymB, but these included a larger proportion of metabolism and regulation genes (**Fig. 7B and S6A**). pSymA dominated every category, but in some categories pSymB-associated families were as widespread across accessory plasmids as pSymA-associated ones (**Fig. S6A**). These included “Posttranslational modification, protein turnover, chaperones,” “Transcription” and “Signal transduction mechanisms.” Gene exchange with accessory plasmids is therefore not limited to pSymA.

Despite this overlap, accessory plasmids carry an extensive repertoire of genes not found in the megaplasmids (**Fig. S7A**). Accessory plasmids harbor frequent recombination, regulation and metabolism functions, and many “Function unknown” or uncategorized genes, marking them as a source of gene novelty (**Fig. 8A**).

**Fig. 8.**
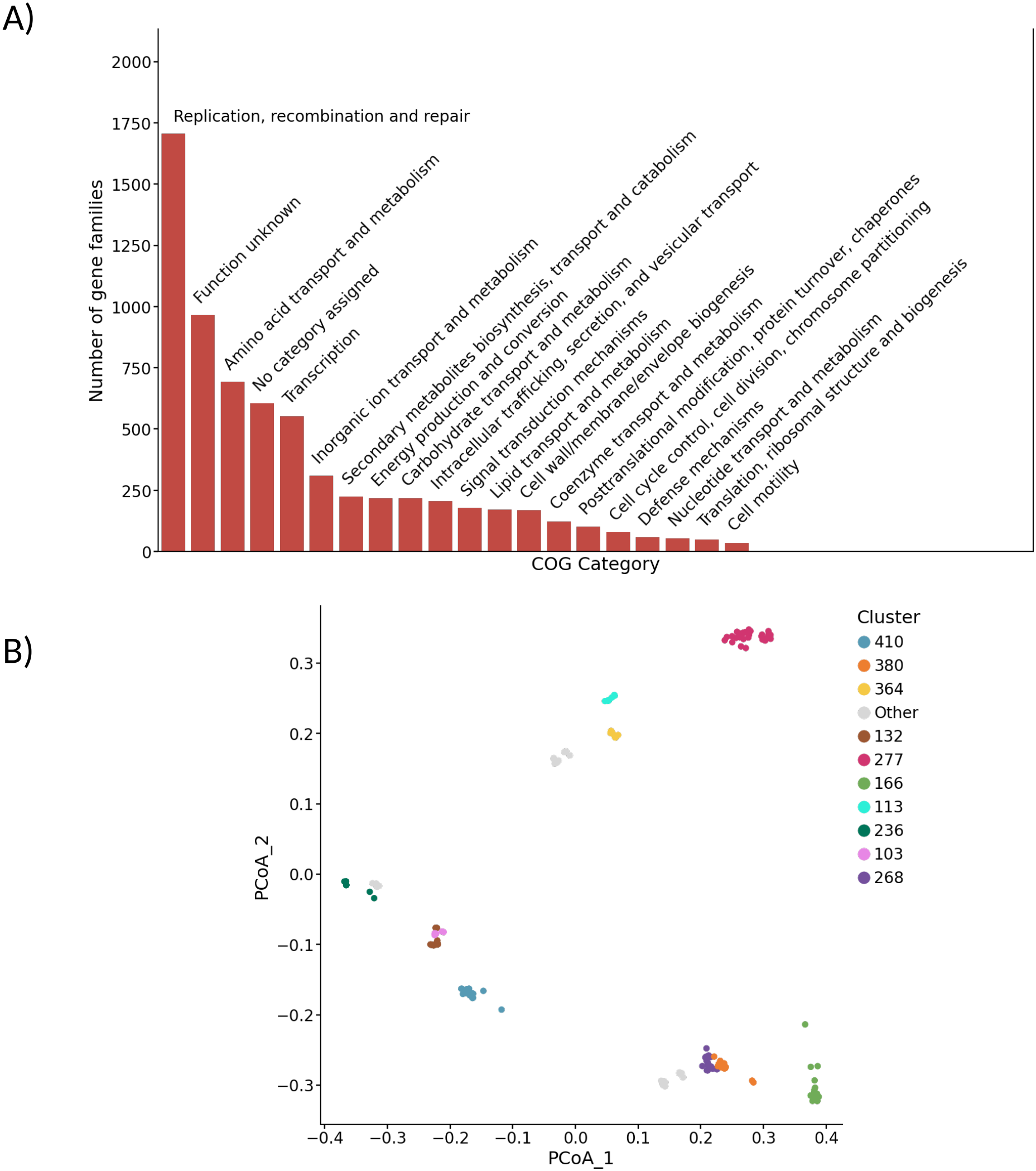
**A)** Number of gene families per functional category (COG) across accessory plasmidome. **B)** PCoA of accessory plasmids based on gene content similarity. PCoA of Jaccard distances computed from COG-annotated orthologous gene presence/absence across accessory plasmids. Points represent individual plasmids, colored by their plasmid cluster (top 10)

Plasmid clusters formed mostly discrete groups in gene-content space, as observed in *Rhizobium leguminosarum* (26), with some showing similarities by proximity, such as clusters 103 and 132 (**Fig. S7B**), likely indicating sharing similar functions or roles (**Fig 8B and S7C**). GOC enrichment suggested differences between clusters: some showed increases in “transcription,” “amino acid transport and metabolism,” or defense mechanisms, others in “intra-cellular trafficking, secretion and vesicular transport” or “nu-cleotide transport and metabolism” (**Fig. S7D and S7E**). Functional studies at the plasmid level are needed to determine whether these reflect specialized roles across niches.

### Accessory Plasmids Encode Biosynthetic and Host-Interaction Functions

Biosynthetic gene clusters (BGCs) were long viewed as static, species-specific features inherited vertically, but reports of plasmid-borne BGCs implicate horizontal transfer in their diversification (99). Large mobilizable plasmids can carry several BGCs (100, 101). These supply secondary metabolites that provide defense, such as antibiotic activity, or competitiveness in stressed environments like nutrient-limited soils or anoxic ecosystems (99, 102).

BGCs were found in 55% of accessory plasmid clusters (81 clusters; 270 plasmids). Unlike on chromosomes and megaplasmids, NRPS clusters predominated, followed by NI-siderophores, terpenes, and homoserine lactones (**Fig. 9A and B and Fig. S8A**). Most of the 472 BGC regions had no characterized counterpart (**Fig. 9A**). Only terpenes, all related to the *S. fredii* NGR234 gibberellin operon (MIBiG BGC0001605), matched a MIBiG reference (56 BGCs, 11.9%). NI-siderophores (95%), RiPP-like clusters (91%), and homoserine lactones (78%) were largely represented in antiSMASH-DB. In contrast, most phosphonates (94%), hydrogen cyanide clusters (93%), and NRPS (74%) fell outside known GCFs. Overall, 52.8% of accessory plas-mid BGCs were neither characterized nor previously observed.

**Fig. 9.**
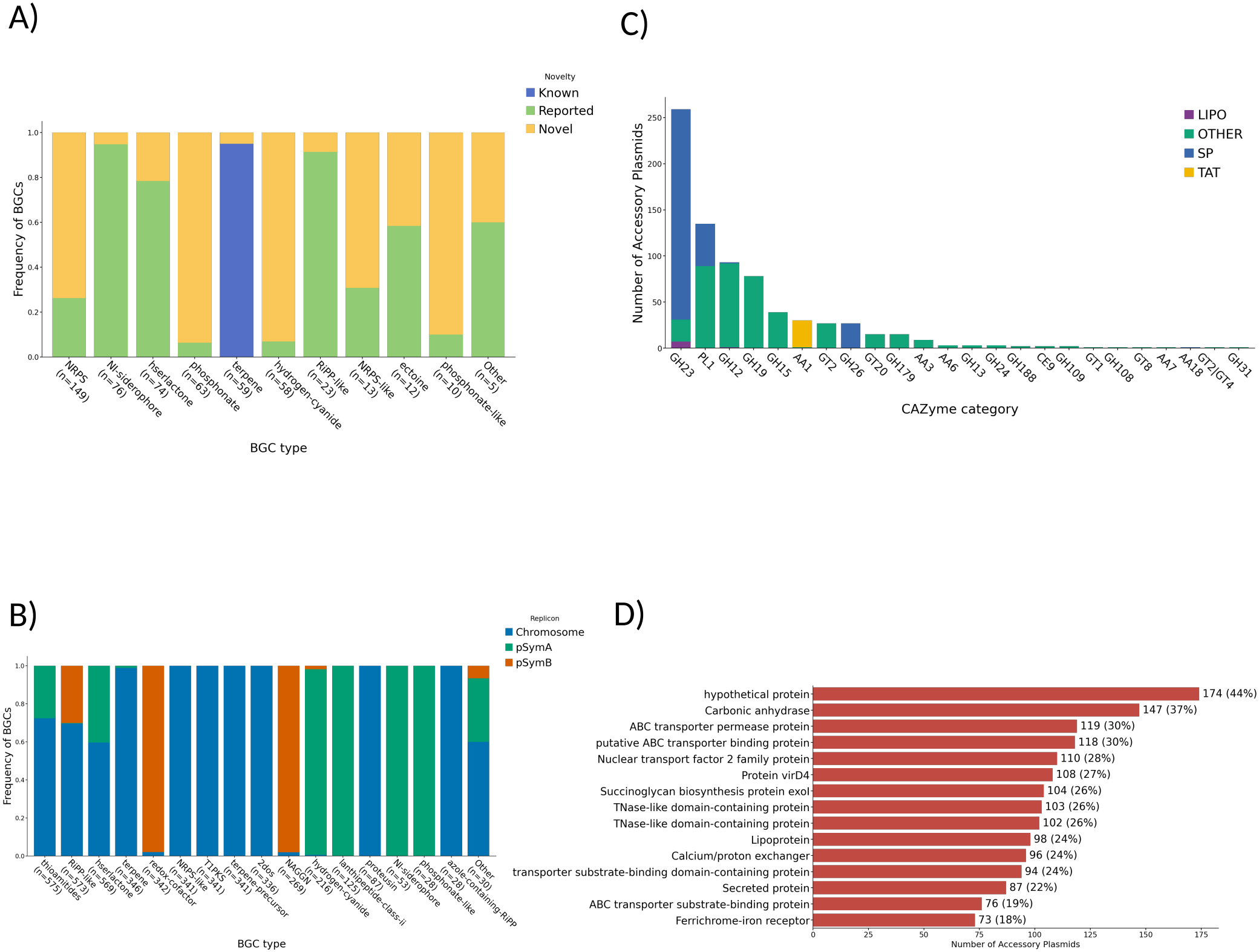
**A)** Novelty classification of accessory plasmid BGCs. Proportion of BGCs of each antiSMASH BGC type that are known (BiG-SCAPE GCF at cutoff 0.3 contains a MIBiG 4.0 reference), reported (within a BiG-SLiCE distance of 0.65 of an antiSMASH-DB v4 GCF), or novel (neither MIBiG association nor within a 0.65 distance from all antiSMASH-DB GCFs). Types with fewer than 10 BGCs are grouped as “Other”. **B)** BGC proportions across chromosomes and megaplasmids. The x-axis shows the antiSMASH BGC types with total numbers. **C)** CAZymes frequency across accessory plasmids, colored by signal peptide. One count for each CAZyme family per plasmid. “Other” means no signal. **D)** Number of proteins with signal peptide across accessory plasmids. One count per plasmid.

Accessory plasmids also encoded CAZymes, including glycoside hydrolases (GHs; most frequent, 10 families), polysaccharide lyases, glycosyltransferases, and auxiliary activity enzymes (**Fig. 9C**). CAZymes contribute to host colonization in phytopathogens (103, 104) and to nodule development. For example, an *S. meliloti lpsB* (GT1) mutation reduced nitrogen fixation efficiency (105), and *Medicago truncatula* GHs degrade Nod factors to levels compatible with infection thread formation (106).

Signal peptide-bearing proteins were encoded in 93% of plasmid clusters (395 accessory plasmids). The most abundant families included substrate-binding proteins, nucleases, lipoproteins, and the succinoglycan biosynthesis protein ExoI (**Fig. 9D**). Secreted nucleases are linked to survival, horizontal gene transfer, and degradation of extracellular DNA barriers during host colonization (107–111), and may similarly aid rhizosphere niche establishment. Deletion of the lipoprotein LppA reduced EPS-I production and competitive colonization (112), and succinoglycan is required for infection thread formation and functional nodules (113, 114). Their co-occurrence suggests that accessory plasmids could modulate EPS-I-dependent symbiosis.

### Horizontal Gene Transfer Events in Accessory Plas-mids Reflect Narrow Host Range

The HGT network was built from 1,779 PLSDB plasmids spanning 61 bacterial genera and 172 species. Plasmid size ranged from 1.7 to 500 kb (median 204 kb). Two lineages dominate the set: Enter-obacterales, led by Escherichia (n = 413), Salmonella (n = 303) and Klebsiella (n = 154); and Rhizobiales, led by Rhi-zobium (n = 260), Agrobacterium (n = 236) and Sinorhizo-bium/Ensifer (n = 109) (**Fig. S9A and S9B**).

Most HGT connections linked our plasmids to plasmids from Hyphomicrobiales (formerly Rhizobiales), and 7 of the 10 most-connected genera belonged to Rhizobiaceae (**Fig. 10A**). This suggests that plasmid exchange is largely restricted to hosts within the family. It is consistent with reports that conjugation is biased toward related hosts and limited by host defense systems and molecular compatibility (115–119).

**Fig. 10.**
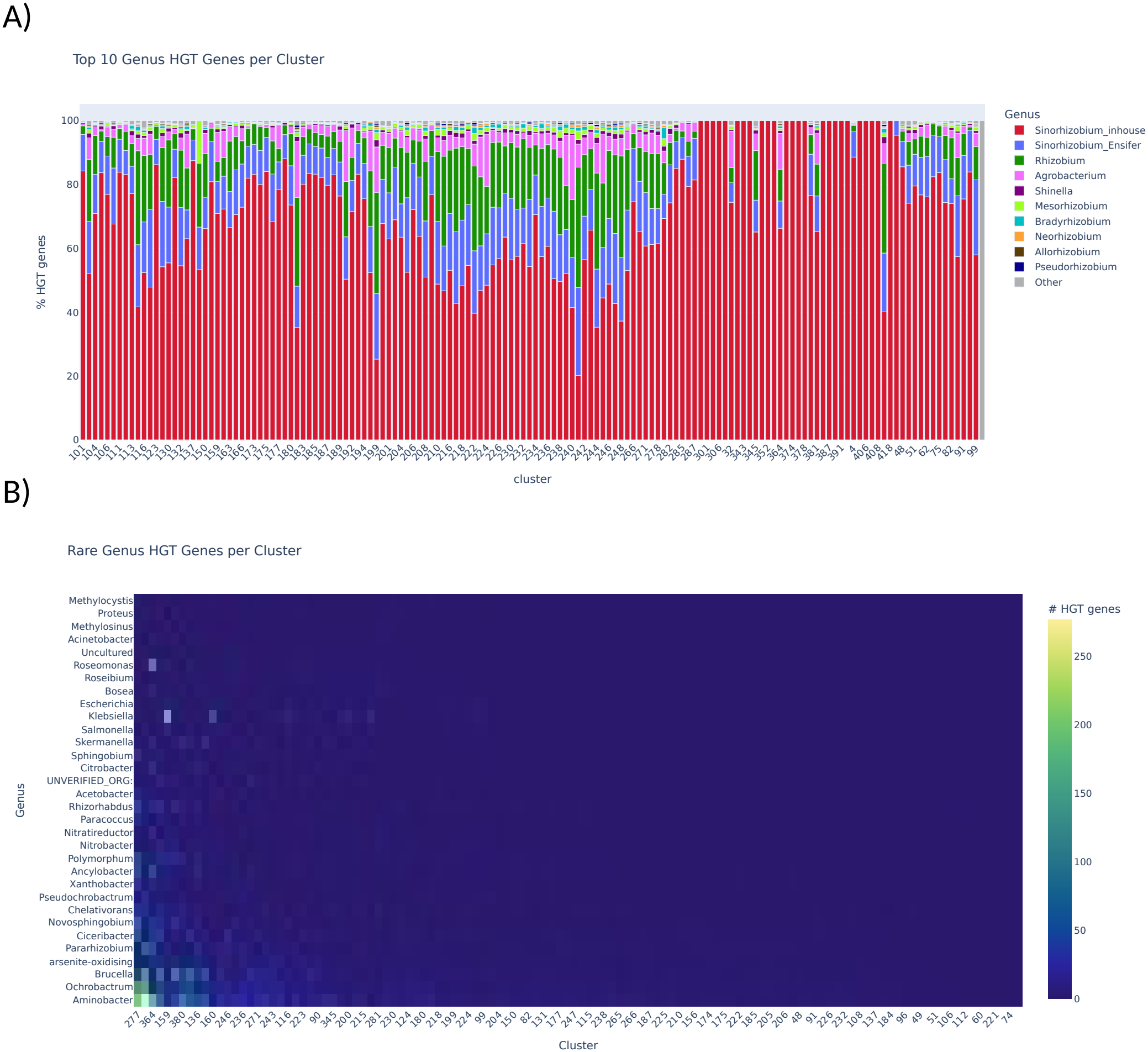
Horizontal gene transfer (HGT) connections between accessory plasmid clusters and other genera. **A)** Stacked bar plot showing the genus composition of HGT connections for each plasmid cluster. The 10 most-connected genera are colored, and all others are grouped as “Other”. **B)** Heatmap of the number of HGT genes shared between plasmid clusters (columns) and genera (rows). Color intensity indicates the number of shared genes. Genera are ordered from top to bottom by increasing number of plasmid clusters to which they are connected.

Connectivity also varied among plasmid clusters, each connecting with a distinct set of taxa (**Fig. S9C and S9D**). This suggests that gene exchange follows lineage-specific routes, which may reflect the ecological history of individual strains. Some plasmids had no connections outside our dataset, suggesting that they circulate only within *S. meliloti* or that their relatives have not yet been sequenced (**Fig. 10A**). Conversely, a few clusters displayed connections with genera that are rare among the plasmids in the reference database (**Fig. 10B**). Accessory plasmids may therefore extend the range of taxa with which *S. meliloti* exchanges genes. Because these inferences depend on the plasmids currently sequenced, additional metagenomic and transcriptomic data is needed.

## Discussion

Despite decades of research on its megaplasmids, the accessory plasmidome of *S. meliloti* has remained largely uncharacterized. Combining long-read sequencing of 346 strains with containment-based de-replication and functional annotation, we recovered 423 large accessory plasmids, present in 76% of strains, most of them (318) absent from PLSDB. These plasmids may contribute to competitiveness in the rhizosphere and during symbiosis.

Accessory plasmids are notably large, with several exceeding 300 kb, a common threshold for a megaplasmid, and the largest (749 kb) approaches megaplasmid size, potentially representing a transitional state between an accessory plas-mid and a megaplasmid. This transition may be promoted by successive fusions with plasmids or other MGEs. Consistent with the framework of (66), accessory plasmids show features of emerging megaplasmids. These include complete replication and partition systems (RepABC), TA systems, and accumulation of host-fitness-related genes through mechanisms such as plasmid fusion and transposons.

Although most accessory plasmids carry the conjugative machinery required for self-transfer, as indicated by their MOB/MPF annotations, this does not necessarily translate into frequent transfer. Limited plasmid mobility has been proposed to reduce gene flow between lineages, thereby allowing their gene content to diverge and accumulate functions shaped by local ecological pressures (19, 26, 66, 120). Consistently, our data fit this pattern. Plasmids shared little sequence beyond backbone regions, and clusters formed discrete groups in gene-content space with distinct functional enrichments. These differences may therefore indicate, at least in part, restricted gene flow among accessory plasmid lineages. Testing this will require transfer experiments such as the mating assays and soil microcosm studies previously performed in *S. meliloti* (16, 64).

Plasmid transfer can be influenced by host-encoded barriers to foreign DNA (121–124). In *S. meliloti*, donor-dependent transformation efficiencies have been observed (11, 27), and this dependence has been linked to the methylation patterns that plasmids acquire in the donor strain. (89, 125). DNA methylation is therefore emerging as an important regulator of HGT in *S. meliloti*. Yet, only a single type I R-M system had been reported, in Rm1021 (88). In contrast to this, we found R-M systems in 153 of 346 genomes, including on 32 accessory plasmids. Interestingly, most of these belong to the type II system. This suggests that R-M is a more widespread barrier to horizontal transfer in this species than previously recognized, and a practical consideration for genetic manipulation, since R-M barriers reduce electroporation efficiency in *S. meliloti* (89, 90, 125). Methylome profiling of plasmids and their hosts will be needed to test this role.

The anti-restriction repertoire was partitioned between replicon classes, with ArdB almost universal on pSymA and ArdC predominant on accessory plasmids, matching the type I and type II systems each is known to counter. This split could reflect timing, with pSymA established before type II systems spread through *S. meliloti* populations, favoring ArdC on later-acquired plasmids. Alternatively, ArdC-carrying plas-mids may have introduced type II systems themselves, since a subset of accessory plasmid clusters encode them. Beyond anti-restriction, accessory plasmids carried diverse defense systems, most commonly DS-20, Gabija, and Lamassu-Fam (126, 127), making them both targets and vectors of host defense. Additionally, *S. meliloti* also harbors a vast variety of TA systems that provide plasmid stability. TA components may serve additional functions; for example, many antitoxins belong to the ribbon-helix-helix (RHH) superfamily, a fold shared with CopG-family regulators that control plas-mid copy number (128, 129), and a CopG was recently identified as important during the symbiosis of *Bradyrhizobium* sp. SUTN9-2 with legumes (130). The occurrence of related gene annotations on *S. meliloti* accessory plasmids makes them candidates for such dual roles.

Accessory plasmids shared about a third (34.6%) of their gene families with the chromosome, pSymA or pSymB, leaving 6,205 gene families found only on accessory plas-mids. Among these, of particular interest are the Biosyn-thetic Gene Clusters (BGC) found in the plasmids, since the HGT of complete BGCs allows the rapid appearance of novel metabolic phenotypes. Our results identify accessory plasmids as a large reservoir of uncharacterized biosyn-thetic potential. More than half of their BGCs had neither a characterized counterpart in MIBiG nor a close relative in antiSMASH-DB. NRPS, the most frequent class, were also largely novel (74%), making them a promising source of new bioactive compounds (131–134) and an interesting source for new biotechnological compounds. NI-siderophores the second most predominant BGC in accessory plasmids, mediate sequestration of insoluble ferric iron, an important trait under alkaline and neutral soil pH conditions where iron bioavailability is limited (135, 136). The only MIBiG match was the gibberellin operon, which interestingly acts as a virulence factor in phytopathogenic bacteria; however, when produced by rhizobia in symbiosis with legumes, it affects nodule size (137), suggesting that accessory plasmids could modulate symbiotic outcomes. BGCs mobilized by accessory plasmids could therefore help *S. meliloti* strains compete in the rhizo-sphere and interact with their host.

Accessory plasmids’ defensive capacity, coupled with lineage-stable inheritance and a narrow host range restricted to the family or even species level, as evidenced by the HGT network, reveals a complex taxonomic landscape for *S. meliloti*. This poses particular challenges for micro-biome–plant interaction studies that link plant traits to genomic variation in associated bacteria, where species definitions may fail to account for the extensively open pangenome characteristic of this species (138)). This accessory plasmid repertoire adds a further layer to gene exchange in *S. meliloti* and may promote the coexistence of genetically distinct sub-populations within the same ecological niche (120). Capturing ecologically and functionally relevant genomic diversity in this species will therefore require plasmid-aware, strain-level frameworks.

Validating these patterns will require wet-lab work: plasmid curing and transfer into plasmid-free backgrounds to test contributions to growth, colonization, and symbiosis; mating assays and soil microcosms to measure actual transfer rates; and methylome profiling with R-M and anti-restriction gene deletions to define their role as barriers to plasmid acquisition.Together, these experiments would establish whether the genomic patterns reported here reflect functional constraints on plasmid acquisition and maintenance in natural *S. meliloti* populations.

## ACKNOWLEDGMENTS

We thank the Sequencing and Discovery Genomics Core of the Stowers Institute for Medical Research for whole-genome sequencing of the MAG strains, the Stowers Computational Core for support during bioinformatic analysis, and the Stowers High-Performance Computing team for computing resources and technical support.

We thank the Genomics Core Facility of Huck Institutes of the Life Sciences (RRID: 365 SCR_023645) at Pennsylvania State University for performing the whole-genome sequencing of the AVT strains.

## DATA AVAILABILITY STATEMENT

Genome assemblies, accessory plasmid sequences, annotations and associated metadata are available on the Stowers Original Data Repository at https://www.stowers.org/research/publications/libpb-2666 upon publication. Raw sequencing reads and genome assemblies have been submitted to NCBI under BioProject PR-JNA1558988 and will be publicly released upon publication. Analysis code is available at https://github.com/juanjo255/AccPlasPipeline.git (archived at https://doi.org/10.5281/zenodo.23816857). MAG strains collection metadata is available in Riley et al. (31).

## Supplemental material

Supplemental material (Fig. S1 to S9 with legends; Supplementary Methods) is provided as a separate file (supplementary.tex).

#

#

