## Supplemental Methods and Figures for "Beyond the tripartite genome: a widespread, mobile and gene-rich accessory plasmidome expands the genomic landscape of the model symbiont *Sinorhizobium meliloti*"

### Deeper details of the methods

### **S1.1 Library Preparation and Sequencing of MAG Strains**

MAG strains. Genomic DNA integrity and concentration were assessed with a TapeStation 4200 (Agilent Technologies, Santa Clara, CA, USA) and a Qubit 2.0 Fluorometer (Thermo Fisher Scientific, Waltham, MA, USA). Libraries were prepared from 250 ng of DNA per sample following the WGS PCR-free protocol of the LongPlex Multiplexing Kit (seqWell, Beverly, MA, USA) and the SMRTbell Prep Kit 3.0 (PacBio, Menlo Park, CA, USA) according to the manufacturers’ protocols (Pacific Biosciences, 2025), with reagent volumes halved except those of the indexed adapters and ligation enhancer. Size selection used a PippinHT (Sage Science, Beverly, MA, USA). Libraries were sequenced with SPRQ chemistry and 30-h movies (ICS v13.3.0.253824; SMRT Link v25.1.0.257715). Because size selection depletes short fragments, the read depth, and therefore the estimated copy number, of small plasmids may be underestimated.

TG strains. Genomic DNA was isolated with the Bacterial Genomic DNA Isolation 96-Well Kit (Norgen Biotek, Thorold, ON, Canada). Samples with low DNA concentration were re-extracted with the DNeasy UltraClean Microbial Kit (Qiagen, Hilden, Germany) or purified with a column-based DNA cleanup kit (Qiagen). Libraries were prepared by tagmentation, which fragments and barcodes DNA in a sequence-independent manner.

### **S1.2 Genome Assembly and Contig Classification**

fastp-long was run with default parameters. Flye v2.9.6-b1802 was run with --pacbio-hifi for HiFi reads and --nano-hq for ONT reads, and ONT assemblies were polished with medaka_consensus using model r1041_e82_400bps_sup_v5.2.0. CheckM completeness used the Ensifer (UID3566) marker set. Contigs were classified by alignment with minimap2 v2.28-r1209 to GCF_000006965.1 (Rm1021). Assembly figures were generated with the cnsplots Python package ([Rashidi, 2026](https://github.com/faridrashidi/cnsplots)).

### **S1.3 Accessory Plasmid Identification, Typing and Copy Number**

All contigs < 1 Mb were also classified with geNomad v1.12 using default thresholds; no filter was applied. PLSDB records were selected when *S. meliloti* was their nearest Mash neighbour in the PLSDB typing annotations according to MOB-typer ([Ondov *et al.,* 2016](https://doi.org/10.1186/s13059-016-0997-x)) or the species named in the NCBI record description. No length filter was applied to PLSDB records; clusters containing megaplasmid-sized sequences such as pSymA or pSymB were removed by the cluster length rule below.

MobMess was run with default parameters. Circularity was taken from the Flye assembly information for study contigs and from the NCBI topology field for PLSDB records. Backbone clusters were retained if all members were < 1 Mb; compound clusters additionally required at least one circular member; and maximal clusters outside plasmid systems additionally required at least one study sequence. Clusters made only of PLSDB sequences were therefore retained only when MobMess placed them in plasmid systems (backbone or compound).

The relaxase criterion (geNomad) applied only to study sequences, and MOB-typer results were not used for classification. Study plasmids < 50 kb with a read depth < 0.5× that of the chromosome were flagged for manual review, as low depth may indicate contamination, plasmid loss or a mixed culture.

3-mer frequencies were computed with Kmertools v0.2.1 (comp oligo -k 3), and Bray–Curtis distances with SciPy pdist ([Virtanen *et al.,* 2020](https://doi.org/10.1038/s41592-019-0686-2)). AF was calculated relative to the longer sequence of each pair.

RepA, RepB and RepC were taken from Bakta replication and partition annotations whose product matched these names. From each RepA, a 4 kb window downstream (plus strand) or upstream (minus strand) was scanned, and a window containing RepA, RepB, and RepC was considered a complete operon. When no complete operon was found, RepA was used only if the plasmid carried a single annotated copy. Representative RepA proteins and the Rh references were clustered with MMseqs2 v18.8cc5c (easy-cluster --min-seq-id 0.9 -c 0.7). Clusters containing a reference took its Rh name; the remaining clusters were named Si01–Si13 in decreasing order of size.

For copy number computation, reads were mapped with minimap2 (-ax map-hifi or -ax map-ont according to read type, --secondary=no), and depth was computed with samtools depth -a. The first and last 1 kb of each contig were excluded before taking medians.

### **S1.4 Functional Annotation**

Bakta was run with the full database. TA hits to domains not specific to TA systems (partitioning protein RepB, ParB-like nucleases, ArdC, HTH regulators) were excluded. Each remaining hit was assigned a Pfam 35.0 family by matching its TASmania description to Pfam family descriptions; descriptions shared by several families (PIN domain, ribbon–helix–helix domain, HicB-like antitoxin) were kept as single groups.

BiG-SCAPE was run with --mix --classify class --alignment-mode local --hybrids-off --include-singletons against Pfam 35.0 and MIBiG 4.0. BiG-SLiCE GCFs were defined at a clustering threshold of 0.65, and BGCs were queried (bigslice --query --n_ranks 3) against the precomputed BiG-SLiCE v2 GCF models of antiSMASH-DB v4.0b2. The novelty of each GCF was expressed as the mean distance of its members to the nearest antiSMASH-DB GCF. GCF networks were visualized with Gephi.

### **S1.5 Gene Content Analysis**

Panaroo was run with --clean-mode moderate --remove-invalid-genes, and eggNOG-mapper in MMseqs2 mode (-m mmseqs) against eggNOG database v5.0.2. COG enrichment was computed for each plasmid cluster against all others and visualized as a dot plot of log2 fold change versus −log10 adjusted P. Gene-content profiles were compared with Jaccard distances on COG-annotated orthologous genes and projected by PCoA and MDS.

### **S1.6 Horizontal Gene Transfer Events**

PLSDB plasmids ≤ 500 kb were aligned to the study plasmids with minimap2 v2.28-r1209, and those sharing ≥ 500 bp of alignment with at least one study plasmid were retained. All-vs-all protein comparisons were performed with MMseqs2 v18.8cc5c (e-value < 10⁻⁴, ≥ 35% identity, ≥ 50% coverage). For each plasmid pair, wGRR was calculated as described by[Pfeifer et al. (2024)](https://doi.org/10.1038/s41467-024-45757-3): the sum of bidirectional best-hit identities divided by the number of proteins in the smaller plasmid (range 0–1). The network was built with NetworkX in Python v3.13 and visualized in Gephi v0.11.3 using the Fruchterman–Reingold layout.

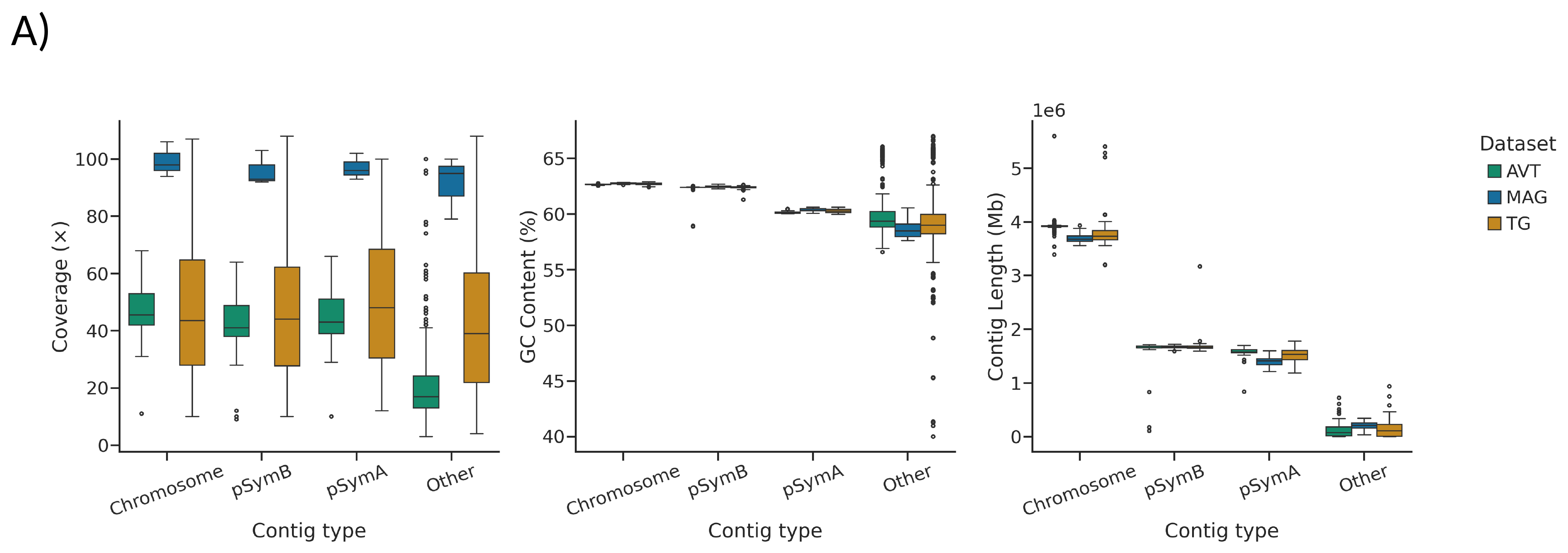

**Figure S1. A)** Left to right: read coverage, GC content, length (Mb), and length per replicon. Other means anything that was not labeled as Chromosome, pSymA, or pSymB.

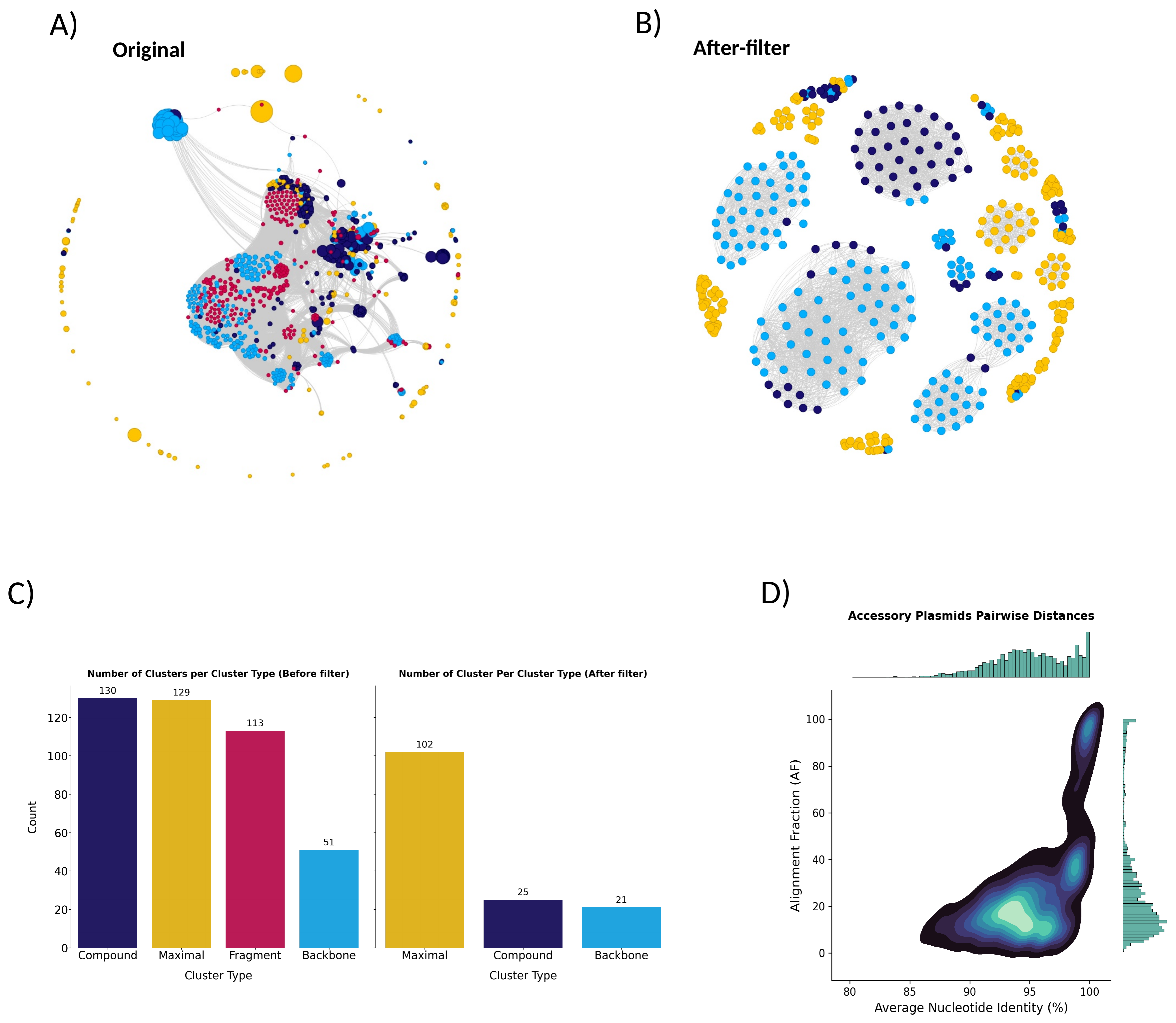

**Figure S2. A)** Containment network of all sequences less than 1Mb inferred by MobMess prior to filtering. Nodes represent sequences; an edge from node i to node j indicates that i is contained within j at ≥ 90% coverage and ≥ 90% identity. Nodes are colored by MobMess cluster type. Compare Figure S2B for the network after filtering (see methods). **B)** Same as A after filtering. Node color indicates category: backbone (sky blue), maximal not in system (amber), fragment (crimson-pink) and compound (deep indigo). **C)** Number of clusters per cluster type in A and B. After filter means after cluster filtering (see supplementary methods) and re-running MobMess classification. **D)** density plot from the all-vs-all average nucleotide identity computation between the accessory plasmids. The x-axis is ANI and y-axis the alignment fraction (AF) computed by skani. Marginal histograms show the distributions of ANI (top) and AF (right). AF belongs to the longest sequence in each pairwise comparison.

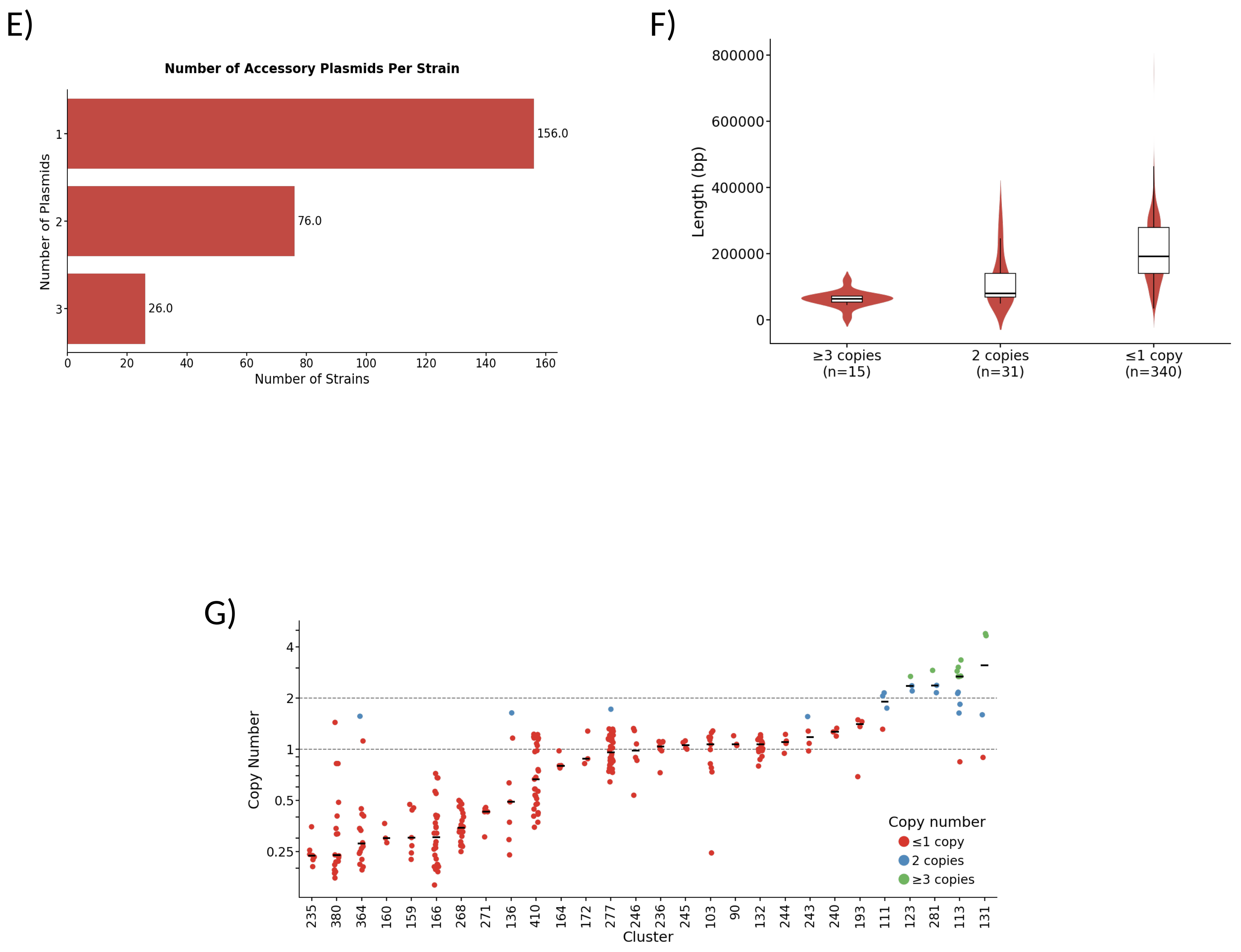

**Figure S2 (continued). E)** Number of strains carrying 1 or more accessory plasmids. **F)** Accessory plasmid length versus copy number. Most accessory plasmids are 1 copy and tend to be large (>100kb). **G)** Strip plot of copy number for each accessory plasmid cluster (clusters ≥3). Some clusters keep copy numbers below 1, possibly due to either sequencing coverage, plasmid loss in isolates, or subpopulations.

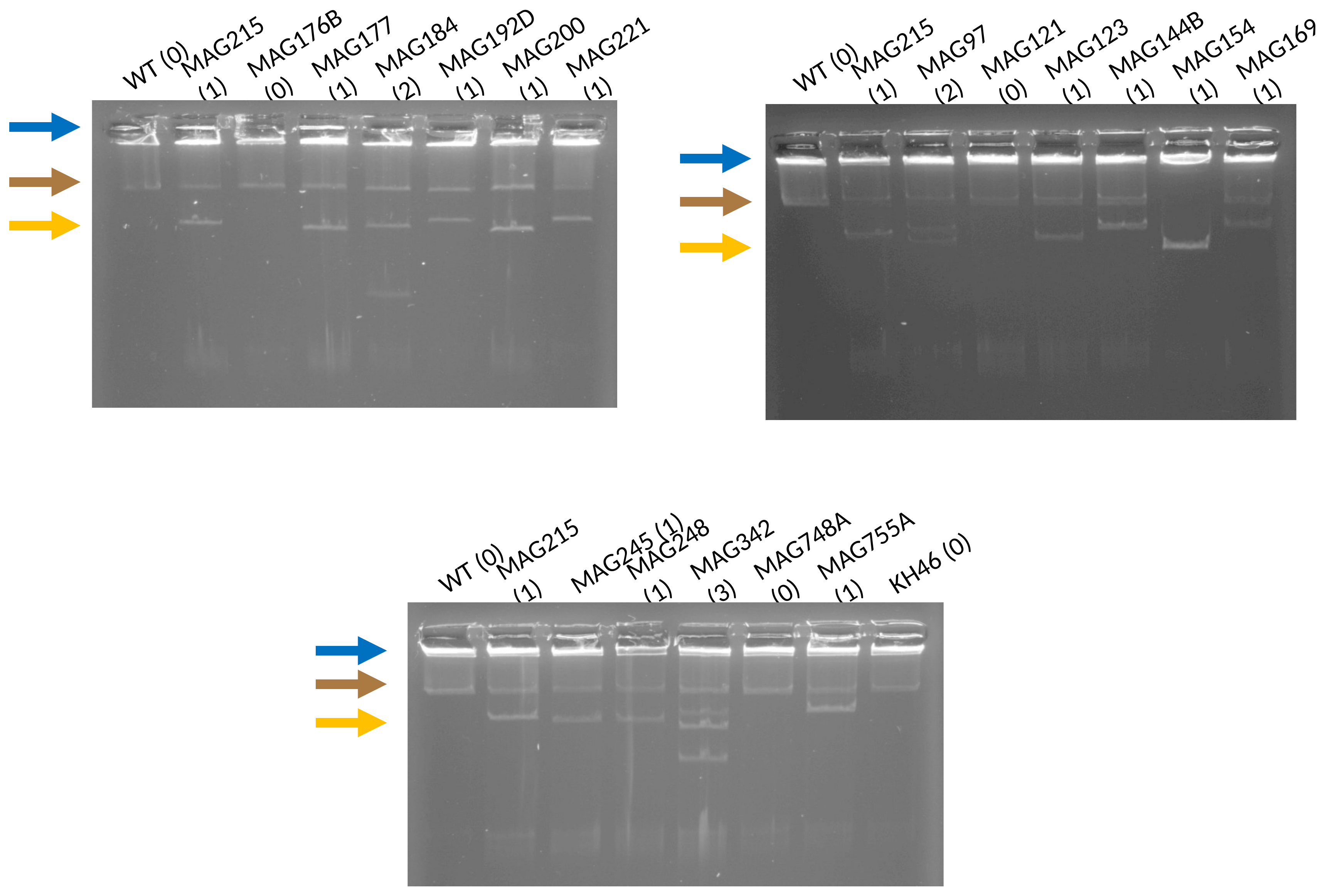

**Figure S2 (continued). H)** Representative agarose gels of plasmid DNA from MAG strains, separated with a modified Eckhardt protocol (65). Numbers in parentheses after each strain name are the number of accessory plasmids. Arrows mark the chromosome (blue), the pSymA and pSymB megaplasmids (brown), and an example accessory plasmid band (yellow).

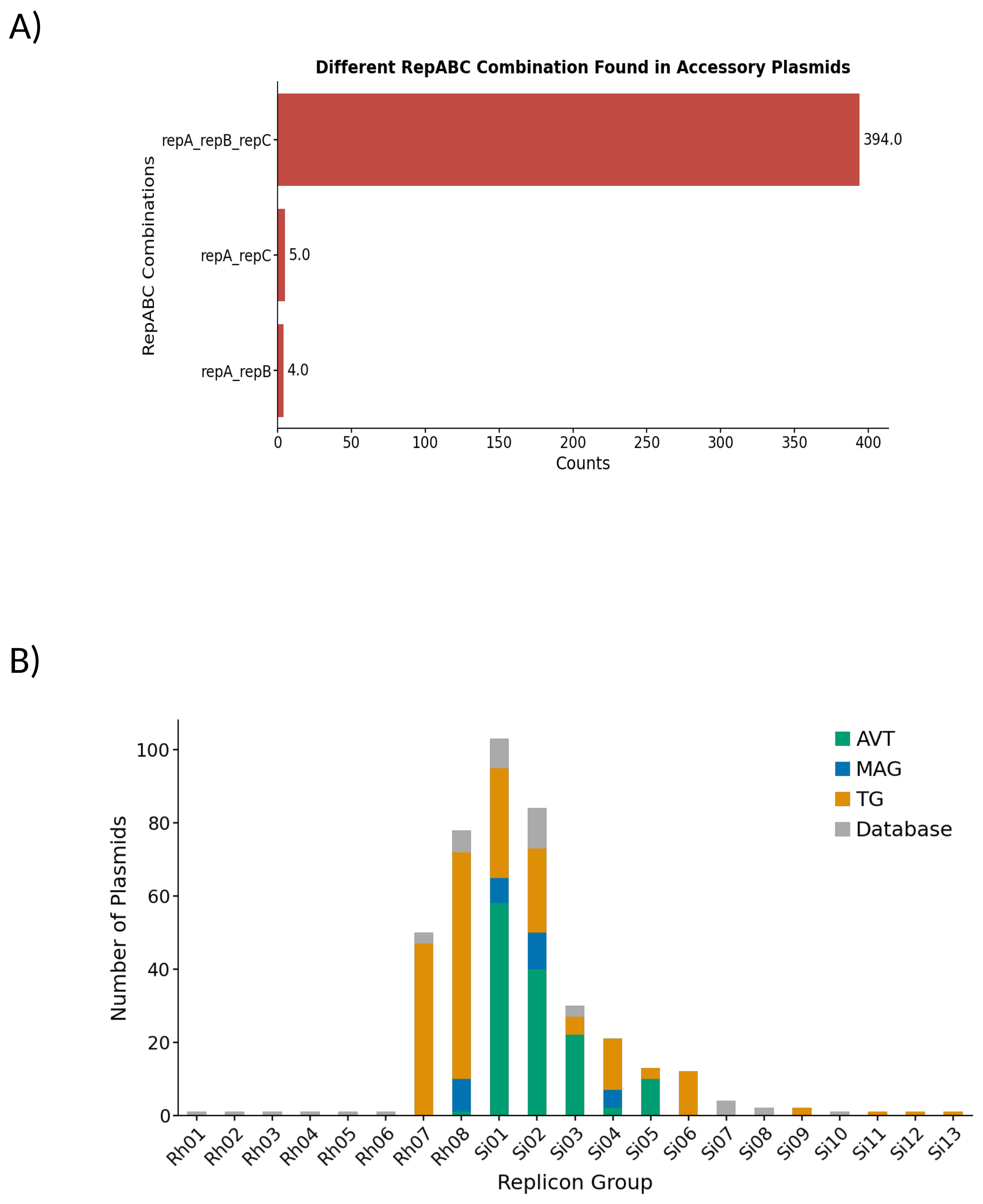

**Figure S3. A)** Different combinations of replication proteins found in accessory plasmids. Most plasmids presented a correctly annotated RepABC. **B)** Distribution of accessory plasmids across replicon groups by dataset origin. The x-axis shows replicon groups, and the y-axis shows the number of plasmids, colored by dataset.

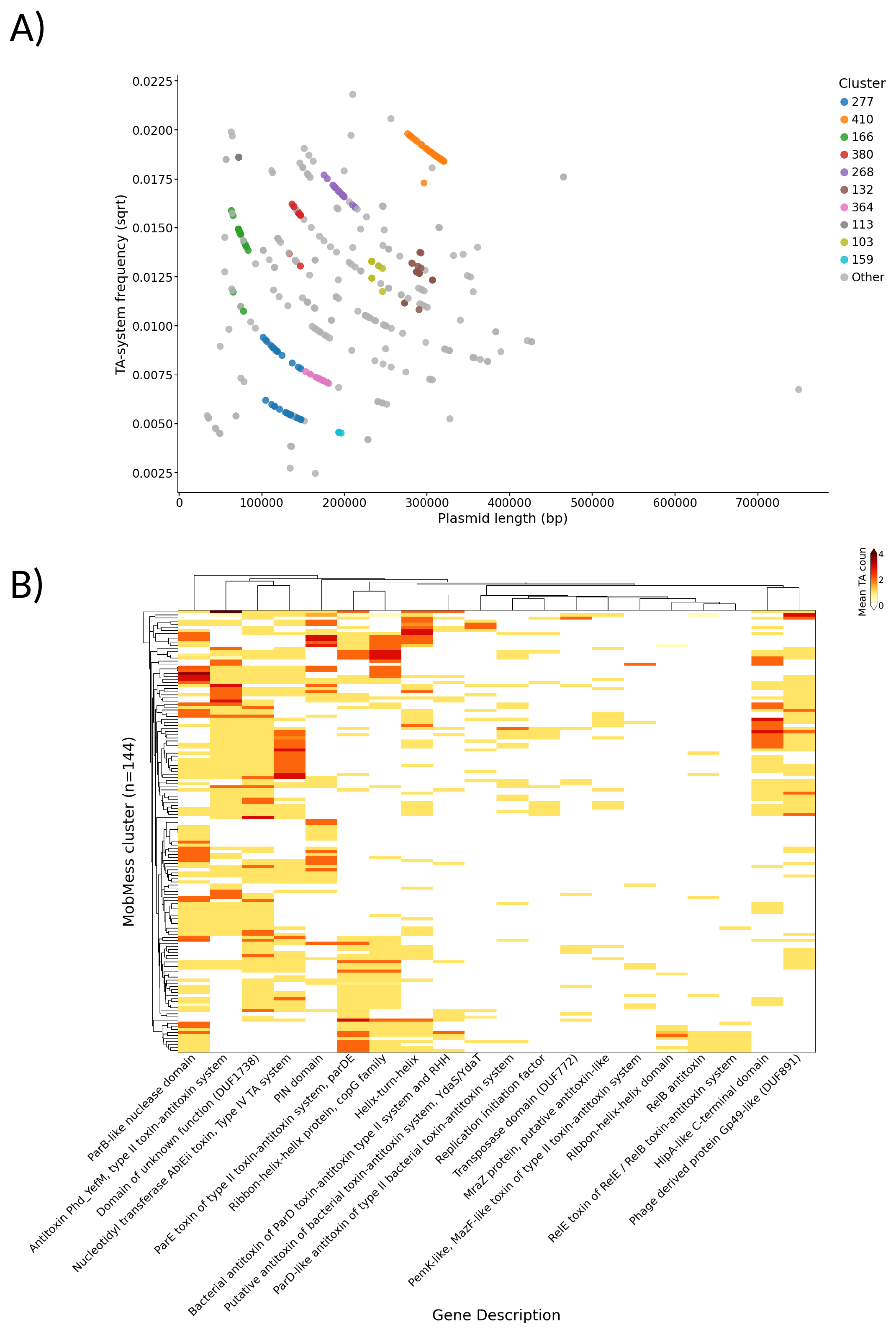

**Figure S4. A)** TA system frequency versus accessory plasmid length across accessory plasmid clusters, as in Bethke et al. (Bethke et al. 2024). Each dot is an accessory plasmid. Top 10 largest clusters are colored. **B)** Heatmap showing the count of toxin-antitoxin (TA) system-associated genes per accessory plasmid cluster using BAKTA annotation. Top 20 most frequent TA-associated gene are plotted. Rows represent accessory plasmids and columns represent TA-associated genes descriptions.

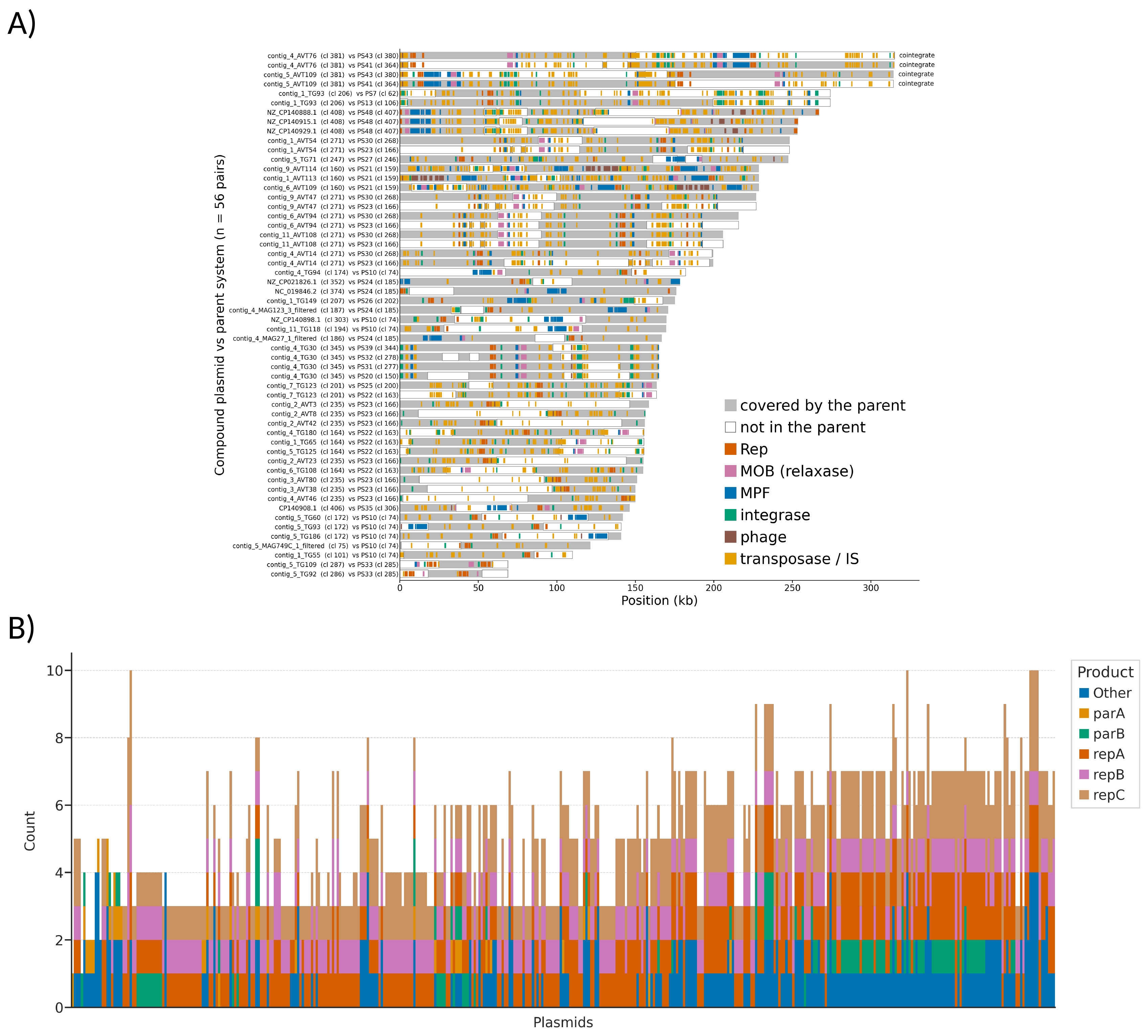

**Figure S5. A)** Sequence gained by compound accessory plasmids relative to their parent backbones. Each row compares one compound plasmid with one of its parent backbones (49 compound–parent pairs; 37 plasmids from 19 MobMess clusters). A compound with several parents appears once per parent. Row labels indicate the plasmid compound ID, the cluster (cl), and the parent system (PS). Rows are sorted by plasmid length. Grey segments show sequence covered by the parent. Outlined white segments show stretches of at least 5 kb absent from that parent; that is, sequence the compound gained relative to it. The legend shows genes related to MGEs along the whole plasmid. **B)** Number of genes related to RepABC per plasmid, sorted left-to-right from smallest to largest. Other represents any annotation related to replication/partitioning.

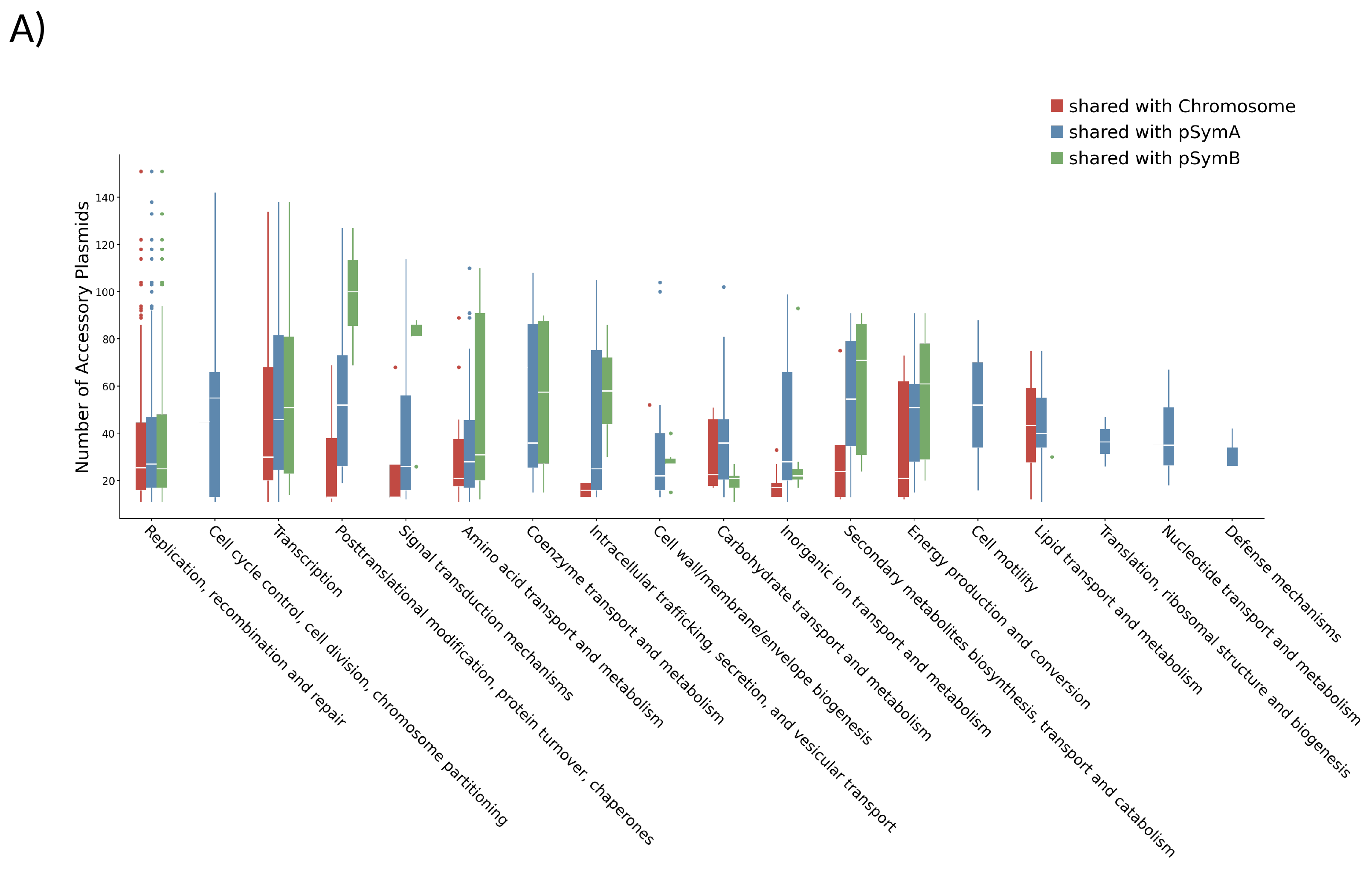

**Figure S6. A)** Distribution of shared COG (gene families) across accessory plasmids. X-axis is the COG categories. Y-axis: the number of plasmids carrying the gene family associated with the COG in x. Y-axis was filtered to values greater than 5.

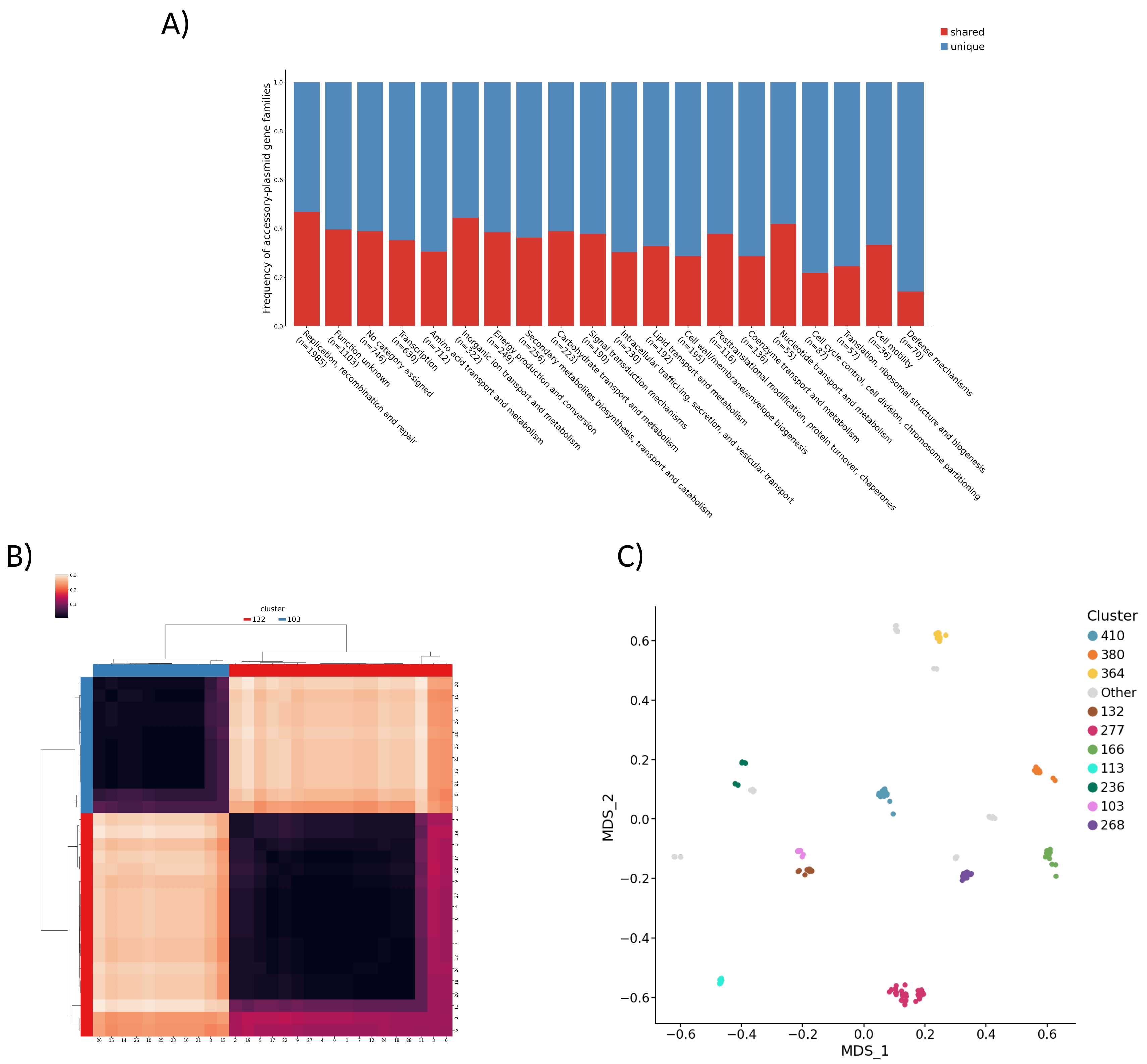

**Figure S7. A)** Proportion of shared versus unique accessory-plasmid gene families across COG functional categories. Bars show the frequency of accessory-plasmid gene families in each COG category that are shared with at least one other replicon (chromosome, pSymA, or pSymB; red) versus unique to accessory plasmids (blue). **B)** Heatmap with Jaccard distances between members of clusters 132 and 103. Rows are colored by the cluster. Each row is a plasmid. **C)** Multidimensional scaling (MDS) projection of accessory plasmids based on COG composition. Computed from COG-annotated orthologous gene presence/absence across accessory plasmids. It allows us to better visualize similarities between high-dimensional data while preserving relative distances. Points represent individual plasmids, colored by their plasmid cluster (top 10).

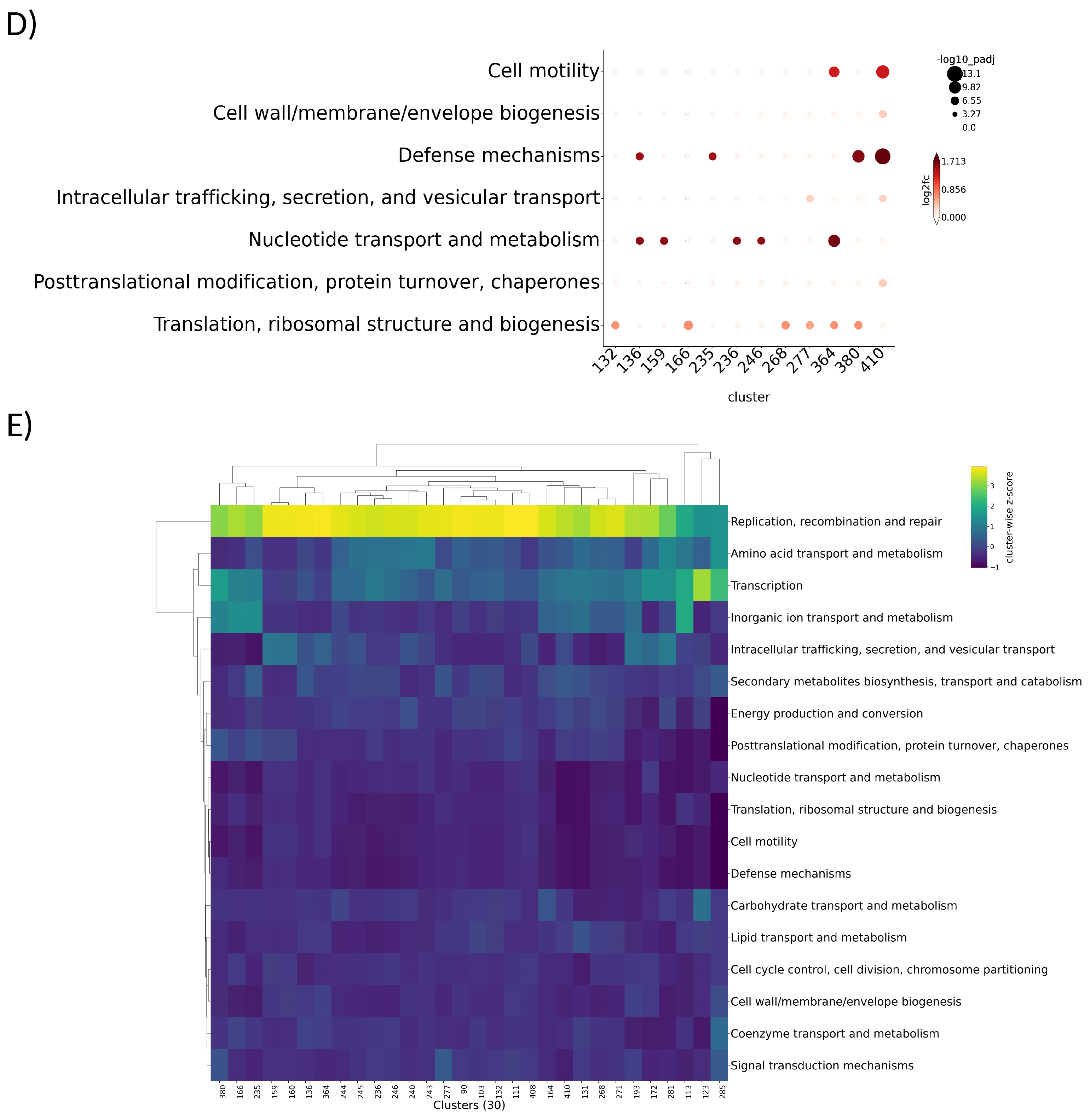

**Figure S7 (continued).** COG composition across accessory plasmids clusters. **D)** COG enrichment across accessory plasmid clusters. Dot plot showing COG category enrichment (rows) for plasmid clusters with significant compositional changes (columns), based on gene annotation (not expression data). Dot color represents log2 fold-change (log2FC) relative to the overall gene content background, and dot size represents statistical significance (-log10-adjusted p-value), determined by Fisher’s exact test with Benjamini-Hochberg correction. Only clusters passing the significance threshold are shown (p>0.05). **E)** Heatmap of COG composition across accessory plasmids clusters. Counts are row-wise z-score. Each row is an accessory plasmid cluster.

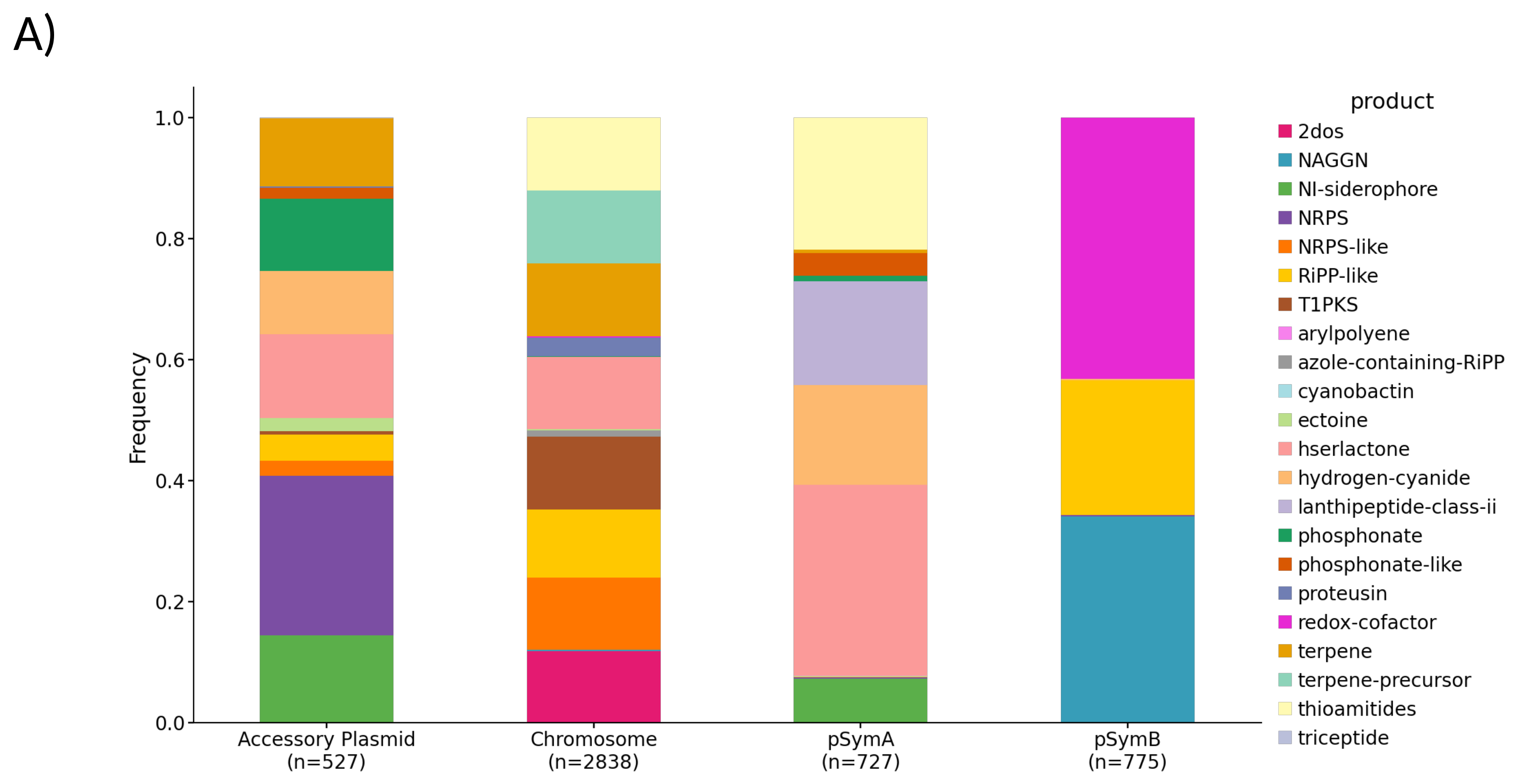

**Figure S8. A)** BGC composition per replicon. One count for each BGC per replicon. Replicons on the x-axis with the total number of BGCs across all genomes. The chromosome concentrates the highest amount and diversity of BGCs, and pSymB the lowest. Accessory plasmids have the highest composition in NRPS

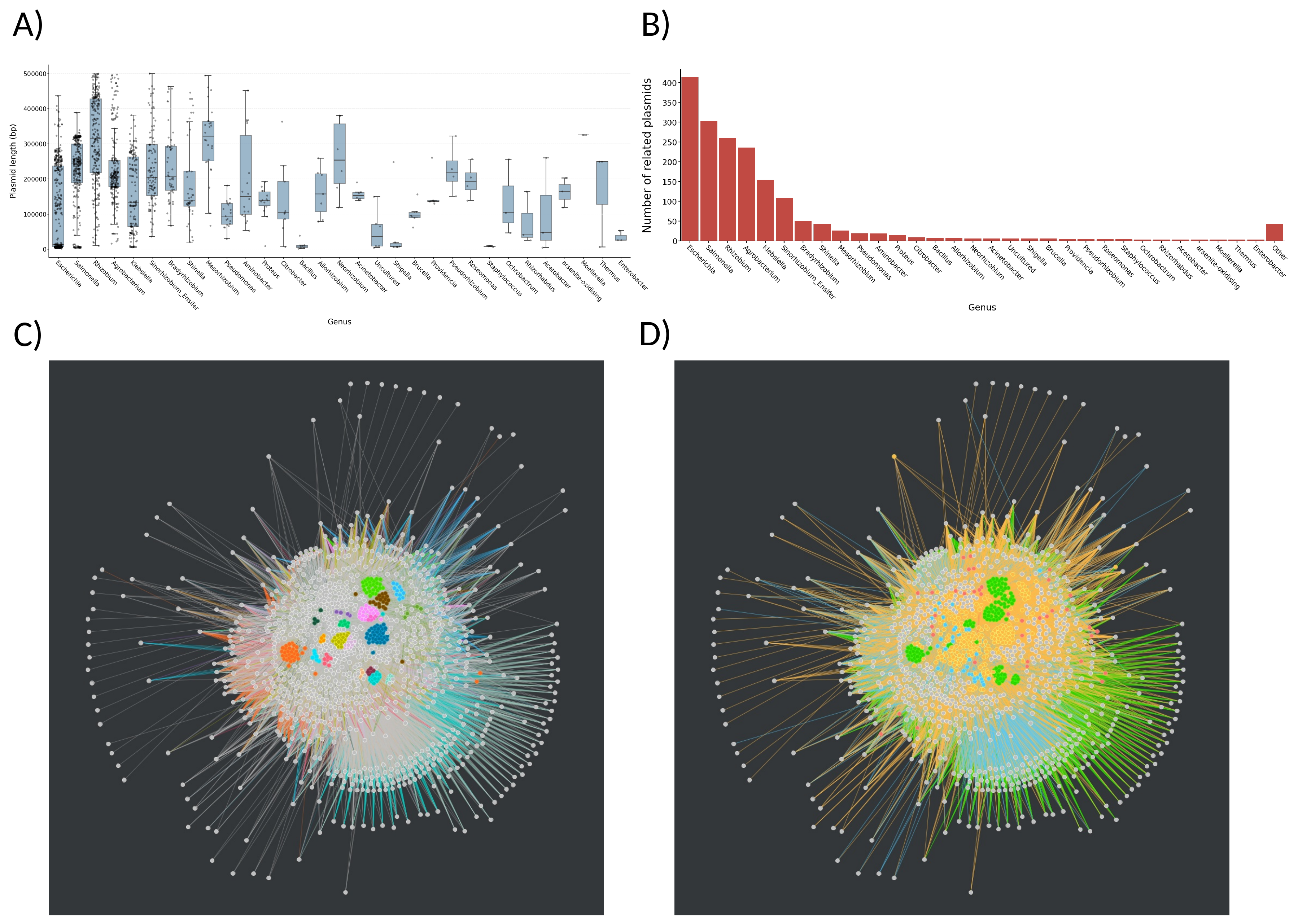

**Figure S9. A)** Boxplot showing length distribution per genus used for HGT analysis. Each dot is a sequence. Only genera with at least 3 sequences are plotted. **B)** Number of plasmids per genus used for HGT analysis. Only genera with at least 3 sequences are plotted. The rest are included in ”Other”. **C)** Horizontal gene transfer (HGT) network colored for the top 20 clusters (> 3 members each). Edges are colored by source and represent wGRR values between 0.001 and 0.1. **D)** Horizontal gene transfer (HGT) network colored by data source. TG (orange), Penn State (green), the reference database (red), and in-house sequencing (blue). Edges are colored by source.
